# Chemical augmentation of the validated HepaRG^TM^ CYP induction test method Part 2: Additional laboratory study supported by mRNA analysis

**DOI:** 10.64898/2026.06.16.732650

**Authors:** Emma Quartermain, Jinkang Zhang, Tim Marczylo, Timothy W Gant, Miriam N Jacobs

## Abstract

Cytochrome P450 (CYP)-mediated biotransformation of endogenous and xenobiotic substances can lead to altered exposure, toxicological impact, or adverse drug reactions. CYP induction data are fundamental to regulatory chemical toxicity hazard assessment because they directly affect the *in vivo* fate of xenobiotics, potentially influencing their safety and efficacy of pharmaceuticals, and impacting the safety assessment of industrial chemicals, and environmental contaminants. Here we report on the third laboratory supplementary validation of an established and previously validated human HepaRG^TM^ *in vitro* method able to detect CYP1A2, CYP2B6, and CYP3A4 induction, to support the expansion of the chemical applicability domain beyond pharmaceuticals. This study was conducted to support the part 1 study with additional robust data.

We established the test method in-house using the 10 previously validated pharmaceutical proficiency chemicals, then tested a further 6 proposed augmentation chemicals, tebuconazole, benfuracarb, chlorpyrifos, N, N-Diethyl-meta-toluamide, fipronil, permethrin, as tested in part 1, and then four additional chemicals: prochloraz, atrazine, pyrimethanil, and chlorpyrifos-methyl. LC-MS/MS was utilised to measure the conversion of a cocktail mixture of prototypical selective CYP probe substrates to their metabolites, in parallel with mRNA measurements.

We achieved high concordance with expected classifications for proficiency and additional chemicals. Comparisons with mRNA-based measurements suggested gene expression may serve as a cost-effective pre-screening tool for CYP1A2 and CYP3A4, though with greater uncertainty for CYP2B6. The data support the robustness of the HepaRG™ method for CYP induction testing and the adoption of the test method in 2026 as an Organisation for Economic Cooperation and Development Test Guideline.

**Plain language summary:** Cytochrome P450 (CYP) enzymes metabolize drugs, pesticides, and other chemicals. Chemicals that increase or decrease CYP enzyme activity can change internal exposure levels, potentially leading to unexpected toxicity or impact drug effectiveness. Reliable *in vitro* methods to assess CYP induction are needed for regulatory chemical safety assessment. This study describes results from a third laboratory applying a previously validated human HepaRG™ cell-based method to assess induction of CYP1A2, CYP2B6, and CYP3A4. After successful in-house implementation using ten reference pharmaceutical compounds, the method was extended to ten more industrial chemicals. CYP induction was evaluated by measuring enzyme activity and changes in gene expression. The test method showed a high level of agreement with expected induction outcomes. Gene expression data supported enzyme activity results, particularly for CYP1A2 and CYP3A4. These results strengthen confidence in the robustness and wider applicability of the method for Organisation for Economic Cooperation and Development Test Guideline adoption.

## 1 Introduction

Cytochrome P450 (CYP) enzymes are central to human Phase I metabolism, controlling the biotransformation of endogenous substances, and xenobiotics including pharmaceuticals, environmental and industrial chemicals. Amongst these enzymes, CYP1A1/1A2, CYP2B6 and CYP3A4 are of particular regulatory importance due to their high hepatic expression and inducibility, their well-characterized regulation by the Aryl hydrocarbon Receptor (AhR), the Constitutive Androstane Receptor (CAR) and the Pregnane X Receptor (PXR) respectively, and their broad substrate selectivity spanning endogenous substances, pharmaceuticals, industrial chemicals, agrochemicals, and environmental contaminants. Activation of these receptors and subsequent CYP induction can significantly alter metabolic clearance rates, modify internal doses, and influence toxicity outcomes. Understanding the induction potential of chemicals on these enzymes is therefore a critical component in mechanistic toxicology and regulatory decision-making. Indeed, historically, one of the most frequently cited limitations of *in vitro* assays concerns the qualitative and quantitative deficiencies in the metabolism of test chemicals, in comparison with *in vivo* assays (Jacobs et al 2013).

In regulatory toxicology, there is a growing requirement for robust, human-relevant *in vitro* systems capable of capturing xenobiotic metabolism, particularly as reliance on animal testing decreases and integrated approaches to testing and assessment (IATAs) using *in vitro* methods develop. The Organisation for Economic Cooperation and Development (OECD) Test Guideline (TG) Programme (TGP) initiated a detailed review on the need for incorporation of metabolism in chemical safety testing, in particular for endocrine disruptors, in 2005, which was later published in 2008 (OECD 2008; Jacobs et al 2008), with a further call for targeted follow-up in 2013 (Jacobs et al 2013). From this work, the cryopreserved, differentiated HepaRG™ cell line, first described in 2002 (Gripon et al., 2002), emerged as a test method solution that should ideally be validated in the longer term, as it is a metabolically competent human *in vitro* hepatic model for CYP induction testing (Jacobs et al 2008, OECD 2008).

The validation and multi-laboratory performance assessment of this test method was reported in 2019 and 2014 (Bernasconi et al., 2019; JRC 2014a, b, c). However, whilst successful, because these validation efforts utilized only pharmaceuticals as proficiency chemicals (as human data were more widely available for pharmaceuticals), some national representatives at the OECD Working Group of National Coordinators of the Test Guidelines Programme (WNT), were concerned that the chemical applicability domain was not sufficiently broad to include industrial chemicals, and so adoption of the first draft Test Guideline (TG) was halted. To address this perceived shortcoming, a targeted weight-of-evidence literature review chemical selection approach was agreed with the WNT (Jacobs et al., 2022) and then implemented. This chemical augmentation list included industrial, pesticidal/agrochemical and food-additive chemicals (Table 1), covering positives (expected inducers) and negatives (expected non-inducers).

**Tab. 1:**
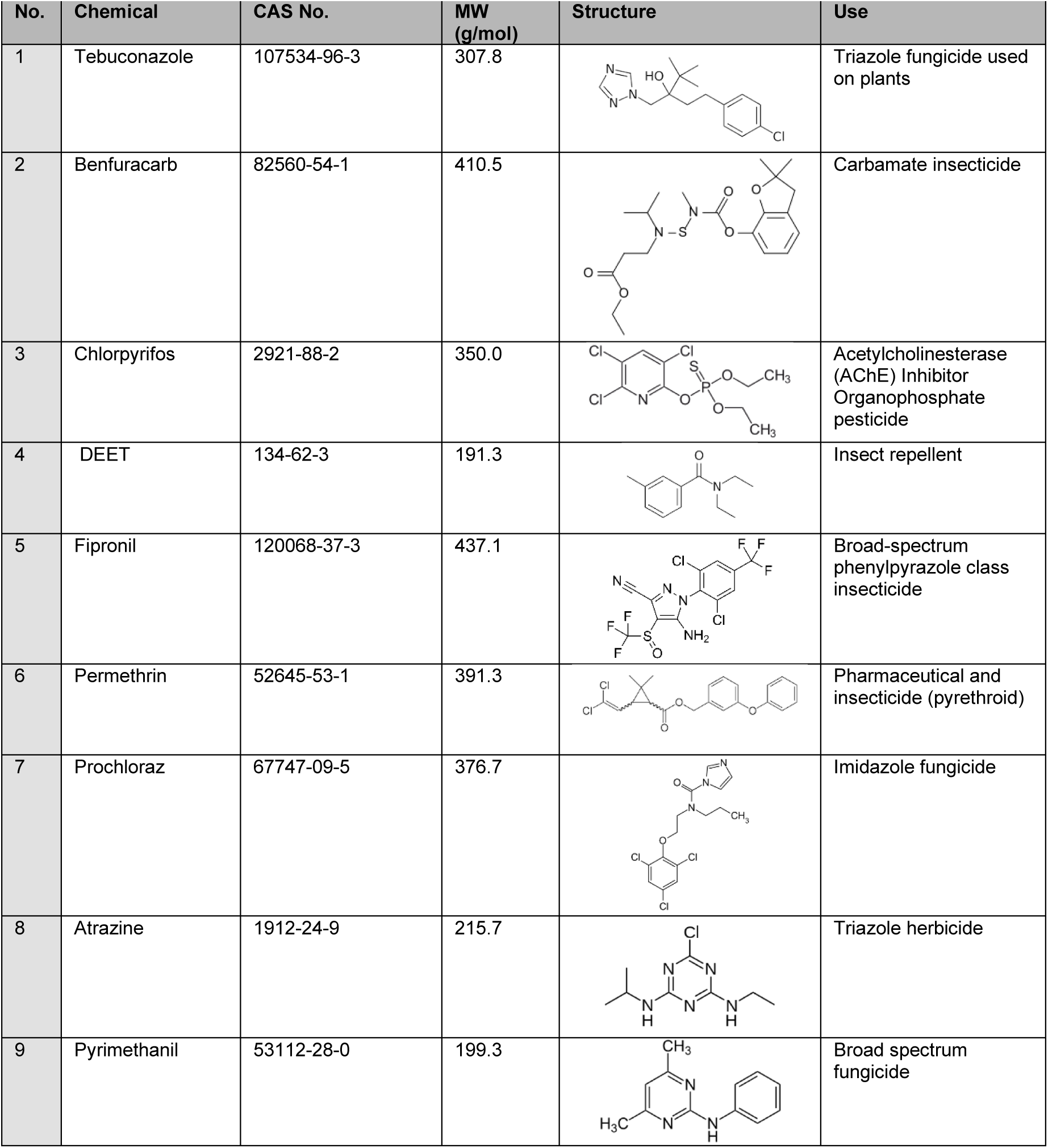
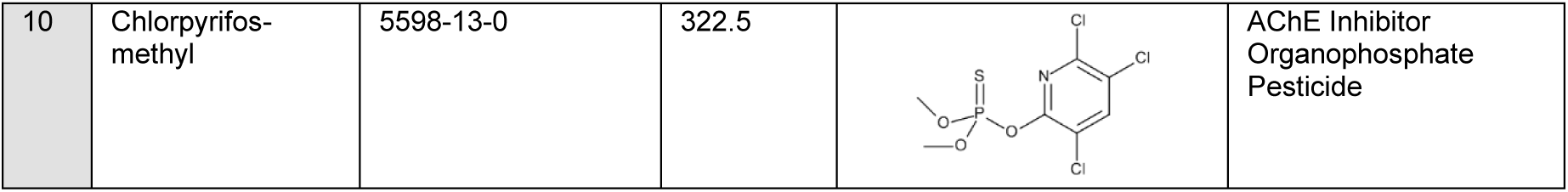
Six Augmentation and Four Additional Chemicals tested in this HepaRG^TM^ CYP induction Test Method Study. OECD TG proficiency chemicals from augmentation (No. 1-6); additional test chemicals (No. 7-10). Chemical name, CAS number, Molecular weight (MW), structure and use are stated.

| No. | Chemical | CAS No. | MW (g/mol) | Structure | Use |
| --- | --- | --- | --- | --- | --- |
| 1 | Tebuconazole | 107534-96-3 | 307.8 |  | Triazole fungicide used on plants |
| 2 | Benfuracarb | 82560-54-1 | 410.5 |  | Carbamate insecticide |
| 3 | Chlorpyrifos | 2921-88-2 | 350.0 |  | Acetylcholinesterase (AChE) Inhibitor<br>Organophosphate pesticide |
| 4 | DEET | 134-62-3 | 191.3 |  | Insect repellent |
| 5 | Fipronil | 120068-37-3 | 437.1 |  | Broad-spectrum phenylpyrazole class insecticide |
| 6 | Permethrin | 52645-53-1 | 391.3 |  | Pharmaceutical and insecticide (pyrethroid) |
| 7 | Prochloraz | 67747-09-5 | 376.7 |  | Imidazole fungicide |
| 8 | Atrazine | 1912-24-9 | 215.7 |  | Triazole herbicide |
| 9 | Pyrimethanil | 53112-28-0 | 199.3 |  | Broad spectrum fungicide |
| 10 | Chlorpyrifos-methyl | 5598-13-0 | 322.5 |  | AChE Inhibitor<br>Organophosphate<br>Pesticide |

Initially, it was agreed with the WNT to establish the test method with 6 proposed augmentation chemicals to be included as proficiency chemicals, in two laboratories, within the Horizon 2020 EU funded GOLIATH project (Legler et al 2021). However, data generated in one of these two laboratories were not sufficiently robust to support the TG adoption (Part 1, Jacobs et al, 2026) and additional funding was sought to conduct supporting validation work in a third laboratory. The data from the third laboratory are reported herein, all conducted in accordance with OECD Good *in vitro* cell culture guidance (OECD 2018a) together with data for an additional four test chemicals to further expand the chemical applicability domain.

Additionally, alongside the established CYP activity endpoint (Bernasconi et al., 2019), parallel CYP gene expression analysis was explored as a more accessible and rapid pre-screen, and to provide additional diagnostic support. Increased CYP enzyme activity often requires both *de novo* protein synthesis from increased gene expression and protein stabilization (Einolf et al., 2014). It also involves allosteric interactions such as heterotrophic effects where a second chemical increases the rate of turnover, such as for example with CYP3A4 and progesterone (Rougee et al 2025). For these reasons changes in gene expression can capture early responses to chemical exposure particularly when these occur through the activation of nuclear receptor pathways leading to increased gene expression mirroring later changes in CYP activity. Gene expression (mRNA) measurements, for CYP1A2, CYP2B6 and CYP3A4, therefore provide a mechanistic measurement that reflects direct transcriptional activation and regulatory receptor engagement (Freeman et al 2007). This makes gene expression profiling a valuable complementary metric particularly when evaluating chemicals with weak induction potential, borderline concentration–response curves, low enzymatic turnover, or substantial between-batch variability.

It is important to note here that only one laboratory conducted the gene expression work, and therefore it has not been shown to be reproducible in another laboratory to date, and it is therefore not included in the OECD TG 445A adopted in 2026 (OECD 2026 in press).

## 2 Materials and Methods

### 2.1 Chemicals and reagents

Dimethyl sulfoxide (DMSO) (D2650, ≥ 99.7%), tebuconazole (107534-96-3, 99.1%), benfuracarb (82560-54-1, 98.3%), chlorpyrifos (2921-88-2, 98.6%), N,N-diethyl-m-toluamide (DEET) (134-62-3, 98.3%), fipronil (120068-37-3, 100%), permethrin (52645-53-1, 100%), prochloraz (67747-09-5, 99.2%), atrazine (1912-24-9, 99%), pyrimethanil (53112-28-0, 98%), chlorpyrifos-methyl (5598-13-0, 99.2%), omeprazole (73590-58-6, ≥ 98.0%), carbamazepine (298-46-4, ≥ 99.0%), phenytoin sodium (630-93-3, ≥ 98.6%), bosentan hydrate (157212-55-0, ≥ 98%), artemisinin (63968-64-9, ≥ 98%), (±)-sotalol hydrochloride (959-24-0, ≥ 98.0%), rifampicin (13292-46-1 ≥ 97%), β−naphthoflavone (6051-87-2, ≥ 98%), phenacetin (62- 44-2, ≥ 98.0%), metoprolol (51384-51-1, 96%), penicillin G sodium (69-57-8, 99.98%), (±)-sulfinpyrazone (57-96-5, ≥ 99%), acetaminophen (103-90-2, 98.0–102.0%), doxorubicin hydrochloride (25316-40-9, 98.0–102.0%), bupropion hydrochloride (31677-93-7, 97.97%), 1’-hydroxymidazolam (59468-90-5, ≥ 97.0%), hydroxybupropion (92264-81-8, ≥ 97.0%); internal standards (+/−) hydroxybupropion−D6 (1184984-06-2, Cat# H−062, 98.84%), acetaminophen-D4 (64315-36-2, Cat# H-909, 99.24%), and α1’−hydroxymidazolam−D4 (Cat# H921, 99.80%), were purchased from Sigma Aldrich (St. Louis, MO, USA). Acetonitrile, methanol and formic acid (Optima; Cat# A117-50) were of LC–MS-grade (Thermo Fisher Scientific) and water (Milli-Q). Phenobarbital sodium salt (50-06-6, 100%) and midazolam hydrochloride (59467-96-8, 100%) were also purchased from Sigma Aldrich with UK Home Office license authorization (due to UK regulatory classification as Schedule III Controlled Drugs).

The proficiency chemicals tested were as previously described in Bernasconi et al., 2019, being omeprazole, carbamazepine, phenytoin sodium, penicillin G sodium, sulfinpyrazone, bosentan hydrate, artemisinin, rifampicin, metoprolol and sotalol hydrochloride. Summary information on these and the additional chemicals selected have been previously published (Jacobs et al., 2022) and are provided in Tab. 1.

### 2.2 Cell Culture

Experiments were carried out using cryopreserved, differentiated HepaRG^TM^ HPR116-TA08 cells (batches #308, #332, #338, #344, #345 and #358), purchased from Biopredic International (Saint Gregoire, France). Cells were stored in liquid nitrogen and were cultured according to providers’ instructions. Medium used was either ‘HepaRG thaw, plate and general-purpose medium with antibiotics’ (basal medium MIL600C + supplement Cat# ADD670C), ‘HepaRG serum-free induction medium with antibiotics’ (basal medium MIL600C + supplement Cat# ADD650C) obtained from Biopredic International, and incubation medium (Williams’ E w/o phenol red supplemented to 25 mM HEPES (Cat# 15630-056) and 2mM L-glutamine (Cat# 25030-024) (Thermo Fisher Scientific)). All media was stored at 4 °C for up to 4 weeks.

The previously described Standard Operating Procedure (SOP) from EURL ECVAM TM2009-14 (JRC., 2009; 2014a; 2018) and Bernasconi et al. (2019) were followed, with additional modifications as outlined in Person et al. (2026) and herein. The revised SOP constitutes the SOP for the adopted TG 445A and is available at https://tsar.jrc.ec.europa.eu/index.php/test-method/tm2009-14.

In brief, HepaRG^TM^ HPR116-TA08 cells were seeded at 100 µL/well (72,000 cells/well) into collagen-I–coated 96-well plates (PLA136) (inner wells seeded; outer wells filled with PBS) and incubated at 37 °C, 5% CO₂, 95% relative humidity. Viability was assessed after thawing by trypan blue exclusion, with an acceptance of 80%. HepaRG thaw, plate and general-purpose medium were renewed six hours after seeding, along with cell morphology observations, and was maintained for at least 65 hours before starting inductions. Cells were treated for 48 hours with reference chemicals (β-naphthoflavone 25µM; phenobarbital 500 µM; rifampicin 10 µM), test chemicals (six concentrations in triplicate; 2 chemicals per plate) and solvent controls (Fig. S1).

### 2.3 Solubility and cytotoxicity assays

Before CYP enzyme induction testing, each of the ten additional chemicals underwent a solubility evaluation in DMSO and serum-free induction medium, followed by cytotoxicity assessments with Alamar Blue reagent according to the validated HepaRG™ CYP induction SOP (TM2009-14). Doxorubicin (8 µM) was used as positive control. This ensured that induction data were generated only under conditions with ≥ 90% cell viability, as required by the test method.

### 2.4 Cytochrome P450 Enzyme Activity Quantification

Enzymatic activity was quantified, against a pre-treatment control, by measuring the metabolic rate of conversion of specific probe substrates to metabolites for CYP1A2 (phenacetin to acetaminophen), CYP2B6 (bupropion to hydroxy-bupropion) and CYP3A4 (midazolam to 1’-hydroxy-midazolam).

#### 2.4.1 LC-MS/MS analysis

LC-MS/MS analysis was carried out on a Thermo Scientific Ultimate 3000 UPLC system coupled with a Q Exactive™ Plus Hybrid Quadrupole-Orbitrap mass spectrometer with ESI (Thermo Fisher Scientific, Bremen, Germany). The LC separation was achieved using a Restek Raptor biphenyl column (100 x 2.1 mm, 2.7 μM, Catalog No. 9309A12) at 30°C with a flow rate of 0.5 mL/min. A gradient elution programme was set for solvent A (water, 2mM ammonium formate with added 0.1% v/v formic acid) and solvent B (80% methanol, 20% acetonitrile, 2mM ammonium formate with added 0.1% formic acid v/v) as 0-0.4 min, 1% B; 0.4-0.5 min, 1-50% B; 0.5-4.5 min, 50-95% B; 4.5-5 min, 95% B; 5-5.01 min, 95-1% B; 5.01-6.5 min, 1% B. The autosampler was maintained at 4°C, and 15 μL of samples were injected from a 96-well collection plate. The mass spectrometer was operated in positive ESI mode with the following parameters: spray voltage = 3.5 kV, capillary temp = 280 *°C,* sheath gas flow rate = 50 arbitrary units, auxiliary gas flow rate = 12 arbitrary units, auxiliary gas heater temp = 380 °C, S-Lens RF level = 60. The MS acquisition was conducted in parallel reaction mode (PRM) (the analytes are shown in Table 2) using a resolution of 17,500, AGC target of 1 x 10^5^ and maximum injection time of 50 ms. All data were acquired in profile mode. Data processing and analysis were conducted using Quan Browser in Xcalibur 4.2 (Thermo Fisher scientific, MA, USA).

**Tab. 2:**
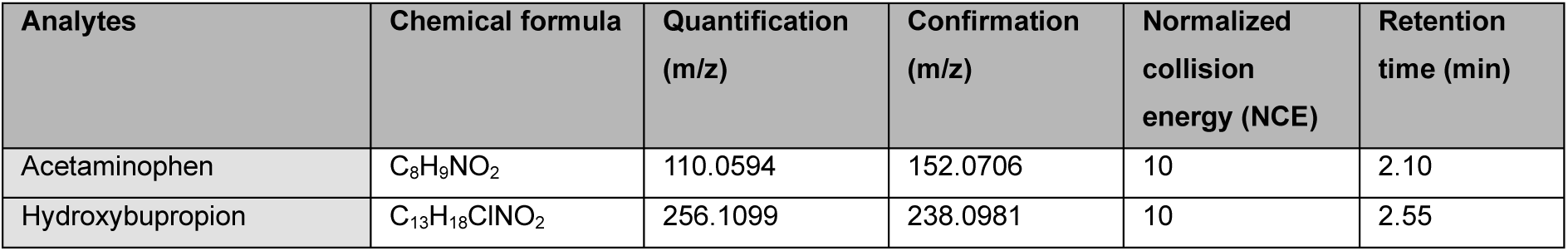
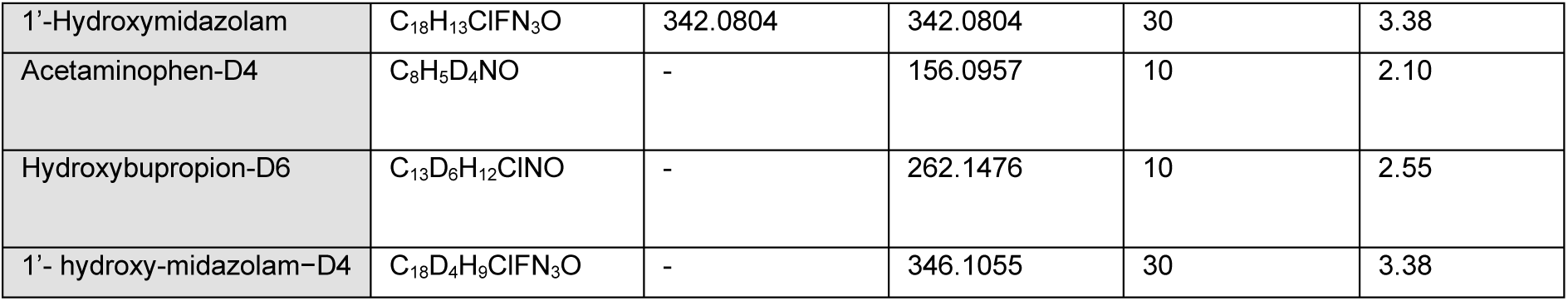
PRM transition parameters for metabolite identification and internal standards.

| Analytes | Chemical formula | Quantification (m/z) | Confirmation (m/z) | Normalized collision energy (NCE) | Retention time (min) |
| --- | --- | --- | --- | --- | --- |
| Acetaminophen | C<sub>8</sub>H<sub>9</sub>NO<sub>2</sub> | 110.0594 | 152.0706 | 10 | 2.10 |
| Hydroxybupropion | C<sub>13</sub>H<sub>18</sub>ClNO<sub>2</sub> | 256.1099 | 238.0981 | 10 | 2.55 |
| 1'-Hydroxymidazolam | C<sub>18</sub>H<sub>13</sub>ClFN<sub>3</sub>O | 342.0804 | 342.0804 | 30 | 3.38 |
| Acetaminophen-D4 | C<sub>8</sub>H<sub>5</sub>D<sub>4</sub>NO | - | 156.0957 | 10 | 2.10 |
| Hydroxybupropion-D6 | C<sub>13</sub>D<sub>6</sub>H<sub>12</sub>ClNO | - | 262.1476 | 10 | 2.55 |
| 1'-hydroxy-midazolam-D4 | C<sub>18</sub>D<sub>4</sub>H<sub>9</sub>ClFN<sub>3</sub>O | - | 346.1055 | 30 | 3.38 |

LLOQ of 5.77 nM for acetaminophen, 3.42 nM for hydroxybupropion and 12.80 nM for 1’-hydroxymidazolam was obtained. Calibration standards and quality controls (Tab. S1 and S2) were used. To correct for any loss of analyte during processing, internal standards of acetaminophen-D4, (±)-hydroxybupropion-D6 and 1’−hydroxy-midazolam−D4 were included.

#### 2.4.2 Protein normalization

For lysis of protein, a modification of the SOP included the replacement of sodium hydroxide (NaOH) with RIPA lysis buffer to ensure no protein degradation, so that gene expression measurement could also be conducted. Determined analytical data for metabolites was normalized for the protein content per well, using the Micro BCA method (Thermo Scientific). The protein concentration of the samples was extrapolated from a bovine serum albumin (BSA) standard curve in 1:10 RIPA buffer using linear regression. Data was only accepted where the coefficient of determination (R^2^) was within threshold (greater or equal to 0.9).

#### 2.4.3 Data processing

CYP induction was determined following protein normalization to ascertain pmol of metabolite formed/min/mg of total protein. Fold changes were calculated relative to the solvent control across three replicates, with each plate including positive controls to confirm assay performance. The n-fold CYP enzyme activity induction was calculated as described in the original SOP (<u>Cytochrome P450 (CYP) enzyme induction in vitro method using cryopreserved differentiated human HepaRG™ cells</u> <u>EURL ECVAM-TSAR</u>; Bernasconi et al 2019; and Jacobs et al 2026 submitted).

Chemicals were characterized based on the following data interpretation procedure (DIP) as recommended by JRC 2014b, c; Jacobs et al submitted, and provided in the OECD TG 445A 2026 (in press):

- “Positive” CYP induction, “positive” CYP induction, if ≥ 2-fold induction is observed from at least two consecutive test concentrations of the 6 test chemical concentrations (with a plausible concentration-response curve).
- “Equivocal” CYP induction, if ≥ 2-fold induction is observed at only one concentration (with a plausible concentration-response curve), or at non-consecutive concentrations.
- “Borderline positive” for CYP induction, if ≥ 2-fold induction is observed at the highest test concentration (or if observed with a non-consecutive response) with a plausible concentration-response curve).
- “Negative” CYP induction in all other cases where ≥ 2-fold induction is not identified.

Further statistical assessment to support results was carried out for each cell batch using a one-way ANOVA compared against the solvent control corrected with Dunnett’s post-hoc test at p > 0.05.

### 2.5 Gene expression (RT-qPCR)

Following metabolite sampling, after removal of the supernatant, the remaining cells in the 96-well plates were homogenized by adding 100 µL/well RLT buffer and RNA was purified using the RNeasy Mini Kit (Qiagen). RNA yield and purity were assessed, and cDNA was synthesized using the High-Capacity RNA-to-cDNA kit (Applied Biosystems). qPCR reactions using 10 ng cDNA, 10 µL SYBR Green Master Mix, 1 µL each of forward and reverse primers (10 µM), and nuclease-free water. Amplifications were performed on a QuantStudio™ 7 Pro (40 cycles) and were quantified by the ΔΔCt method (Livak et al., 2001), normalized to TATA-binding protein (TBP), a previously assessed optimal single reference gene for stability (Ceelan et al., 2011). For benchmarking in the analysis, a 2-fold threshold was set for concordance checks. Primers (5′→3′) were designed via PrimerQuest™ (IDT) (Tab. S3). A one sample t-test was used to compare the mean of all concentrations tested against the solvent control. Statistical significance was defined at p < 0.05.

## 3. Results

### 3.1. Positive control CYP induction

All plates tested included reference chemicals (β-naphthoflavone; CYP1A2, phenobarbital; CYP2B6, rifampicin; CYP3A4), which consistently gave expected induction of relevant CYPs above the acceptable criteria of ≥2-fold induction (Tab. S4A-C).

### 3.2. Establishment with pharmaceutical proficiency chemicals

Expected classifications were seen for all three CYPs, demonstrating successful method transfer. Under the stricter updated DIP recommended by the original ECVAM Scientific Advisory Committee (ESAC) review (JRC., 2014 b, c) and from the part 1 study, several chemicals show as equivocal for CYP1A2/2B6, mirroring earlier observations of greater batch variability for these (Jacobs et al., 2026 submitted) (Tab. 3). Bosentan hydrate leads to a CYP2B6 equivocal classification, yet statistics indicate a reduction in CYP2B6 activity. Batch agreement was ∼70% for CYP1A2/3A4 and 40% for CYP2B6 across experiments when compared with GOLIATH results (Jacobs et al., 2026 submitted) (Tab. S5).

**Tab. 3:**
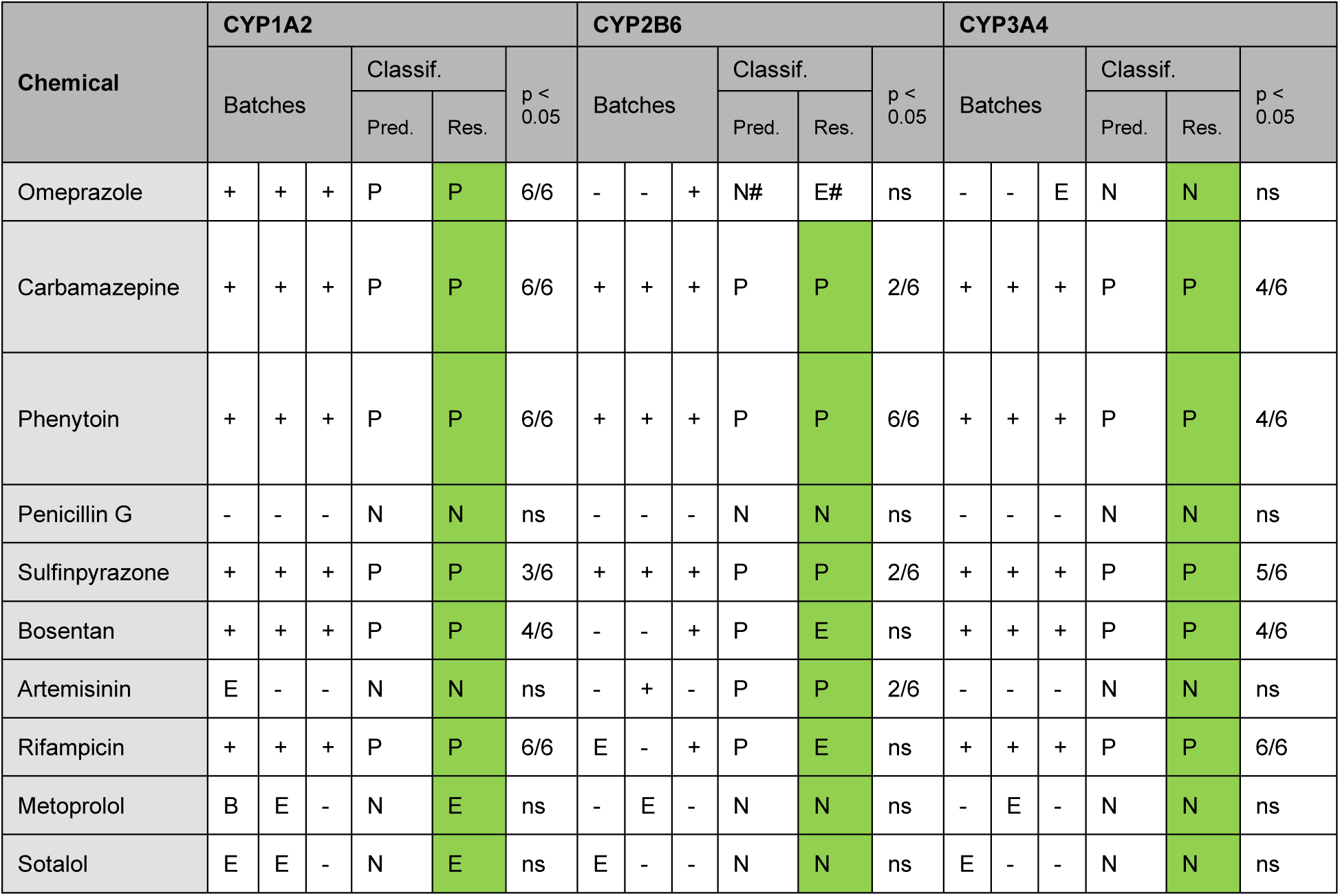
Pharmaceutical proficiency chemicals compared against the expected enzymatic activity. As taken from the original validation study (JRC., 2014a, Bernasconi et al., 2019). Green: agreement. CYP induction per batch (308 / 344 / 345) (+: Positive; −: Negative; E: Equivocal; B: Borderline; P: Inducer; N: Non-inducer; #: Excluded from proficiency testing based on the validation ring trial results (JRC, 2014). Pred.: Predicted classification; Res.: Result from study; Classif: Classification positive/ negative / equivocal. Significance assessed using one-way ANOVA with significant data points of 6 shown (n = 6). See Tab. S5. ns (0/6) = not significant

### 3.3 Solubility and cytotoxicity

For the previously determined six augmentation chemicals, solubility and cytotoxicity testing were carried out in the part 1 GOLIATH study (Jacobs et al., 2026 submitted). The additional four chemicals were initially tested at solubility concentrations of 40 mg/mL for DMSO, and 80 µg/mL, for medium. Prochloraz was only soluble at 20 mg/ml in DMSO and medium. Cytotoxicity assessment was then determined and is provided in Tab. 4.

**Tab. 4:** Solubility and non-cytotoxic concentrations for the four additional tested chemicals.

| Chemical | Solubility in DMSO |  | Solubility in medium |  | Concentrations for CYP induction |  | Final solvent concentration |
| --- | --- | --- | --- | --- | --- | --- | --- |
|  | mg/mL | mM | µg/mL | µM | µg/mL | µM |  |
| Prochloraz | 20 | 53.1 | 20 | 195 | 10.0 – 0.31 | 26.5 – 0.83 | 0.1% DMSO |
| Atrazine | 40 | 185.4 | 80 | 360.9 | 40.0 – 1.25 | 185.4 – 5.80 | 0.1% DMSO |
| Pyrimethanil | 40 | 200.7 | 80 | 401.4 | 40.0 – 1.25 | 200.7 – 6.27 | 0.1% DMSO |
| Chlorpyrifos-methyl | 40 | 124.0 | 80 | 248.1 | 40.0 – 1.25 | 124.0 – 3.86 | 0.1% DMSO |

### 3.4 Augmentation chemicals

Across the augmentation set, we observed robust and largely predicted induction patterns for CYP1A2 and CYP3A4, with more variability for CYP2B6. Tebuconazole and benfuracarb were positive for all three CYPs across batches. DEET and fipronil positively induced CYP1A2, CYP3A4 and (for DEET) CYP2B6. The literature weight of evidence assessment that fipronil induction of CYP2B6 occurred was inconclusive/positive, although one publication did support this induction (Franzosa et al., 2021). The results reported here (and in part 1 Jacobs et al 2026), support a positive conclusion. Chlorpyrifos showed the expected pattern (CYP1A2 positive; CYP2B6 negative), yet CYP3A4 was found to be an equivocal inducer supported by statistically significant induction (p > 0.05) (Tab. S5C). Permethrin was predicted to be a non-inducer for all CYPs, yet CYP1A2 and CYP2B6 were found to be equivocal with no significance suggesting it is a negative. Results are indicated in Tab. 5 (Fig. S2A-F; Tab. S5A-C; S6).

**Tab. 5:** Augmentation chemicals compared against the expected CYP activity (Jacobs et al., 2022). Green: agreement. Grey: inconclusive predictive induction. CYP induction classification per batch (332 / 344 / 345) and overall. (P = Positive; N = Negative; E = equivocal; B = borderline). Pred.: Predicted classification; Res.: Result from study; Classif: classification positive/negative/equivocal. Significance assessed using one-way ANOVA with number of significant data points (n = 6). See Tab. S5A-C. ns (0/6) = not significant.

| Chemical | CYP1A2 |  |  |  |  |  | CYP2B6 |  |  |  |  |  | CYP3A4 |  |  |  |  |  |
| --- | --- | --- | --- | --- | --- | --- | --- | --- | --- | --- | --- | --- | --- | --- | --- | --- | --- | --- |
| | Batches | | | Classif. | | $P < 0.05$ | Batches | | | Classif. | | $P < 0.05$ | Batches | | | Classif. | | $P < 0.05$ |
|  |  |  |  | Pred. | Res. |  |  |  |  | Pred. | Res. |  |  |  |  | Pred. | Res. |  |
| Tebuconazole | + | + | + | P | P | 4/6 | + | + | + | P | P | 5/6 | + | + | + | P | P | 5/6 |
| Benfuracarb | + | + | + | P | P | 3/6 | + | + | + | P | P | 6/6 | + | + | + | P | P | 4/6 |
| Chlorpyrifos | + | + | + | P | P | 2/6 | - | - | - | I | N | ns | - | + | - | P | E | 3/6 |
| DEET | + | + | + | P | P | 5/6 | + | + | + | P | P | 6/6 | + | + | + | P | P | 6/6 |
| Fipronil | + | + | + | P | P | 1/6 | E | B | E | NA | E | ns | + | + | + | P | P | 2/6 |
| Permethrin | - | E | E | N | E | ns | E | - | E | N | E | ns | - | - | E | NA | N | ns |

### 3.5 Prochloraz, Atrazine, Pyrimethanil and Chlorpyrifos-methyl

The additional chemical set of prochloraz, atrazine, pyrimethanil, chlorpyrifos-methyl was intended to expand the chemical applicability domain, in accordance with the modular approach to validation (Hartung et al., 2004), targeting expected isoform-selective responses and weaker/borderline cases.

Prochloraz was a non-inducer for CYP2B6 and an equivocal inducer for CYP3A4, supporting previous data reporting no effect on CYP2B6 and an equivocal inducer for CYP3A4 (Braeuning, Mentz et al., 2020; Schmidt, Lichtenstein et al., 2021). Whilst the chemical selection (Jacobs et al., 2022), considered prochloraz to be inconclusive in relation to CYP1A2 induction, our results support this chemical being classified as a plausible inducer for enzymatic activity and mRNA (Fig. 1A, 3A; Tab. 7). Atrazine was expected to be an enzymatic activity inducer for all CYP isoforms tested (Abass, Lämsä et al., 2012). In contrast, this study identified atrazine as an equivocal inducer for CYP1A2/3A4, with no significance. Further batch testing is therefore needed as recommended, for confirmation. Atrazine was a weak CYP2B6 inducer, but again, additional batch testing would be needed to confirm this finding (Fig. 1B). For atrazine, CYP2B6 and CYP3A4 mRNA were also increased (Fig. 3B; Tab. 7).

**Fig. 1:**
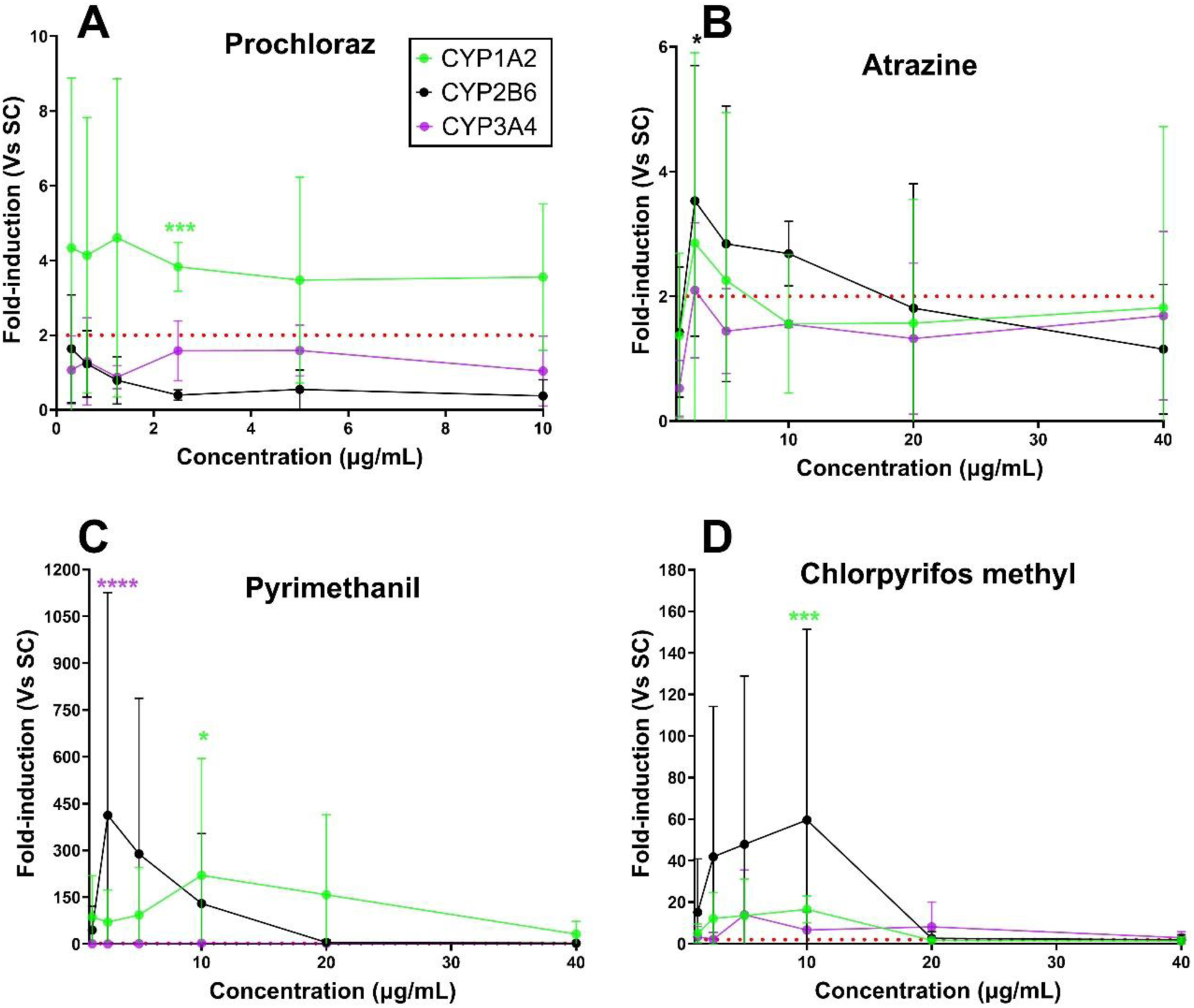
Enzymatic fold change concentration response for four additional chemicals tested. CYP1A2 (green), CYP2B6 (black) and CYP3A4 (purple) are shown. (A) Prochloraz, (B) Atrazine, (C) Pyrimethanil, (D) Chlorpyrifos-methyl. Three independent replicates (n = 3) (mean ± SD). Significance was assessed using one-way ANOVA * = P ≤ 0.05; ** = P ≤ 0.01; *** = P ≤ 0.001; **** = P ≤ 0.0001. GraphPad Prism Software v11.

Pyrimethanil and chlorpyrifos-methyl, activate AhR and are predicted inducers of CYP1A2 (Medjakovic, et al., 2014), and this was supported across batches tested (Tab. 6). Previously lacking data, both are equivocal inducers of CYP2B6 and weak inducers of CYP3A4 (Tab. 6; Fig. 1C-D; S5A-C and S7).

**Tab. 6:** Additional test chemicals compared to the expected CYP activity (Jacobs et al., 2022). Green: agreement. Grey: inconclusive predictive induction. CYP induction classification per batch (332 / 338 / 358) and overall. (P = Positive; N = Negative; E = Equivocal; B = Borderline). Pred.: Predicted classification; Res.: Result from study; Classif: Classification Positive/ Negative / Equivocal. Significance assessed using one-way ANOVA with number of significant data points (n = 6). See Table S5A-C. ns (0/6) = not significant

| Chemical | CYP1A2 |  |  |  |  |  | CYP2B6 |  |  |  |  |  | CYP3A4 |  |  |  |  |  |
| --- | --- | --- | --- | --- | --- | --- | --- | --- | --- | --- | --- | --- | --- | --- | --- | --- | --- | --- |
| | Batches | | | Classif. | | $p < 0.05$ | Batches | | | Classif. | | $p < 0.05$ | Batches | | | Classif. | | $p < 0.05$ |
|  |  |  |  | Pred. | Res. |  |  |  |  | Pred. | Res. |  |  |  |  | Pred. | Res. |  |
| Prochloraz | + | + | B | NA | P | 1/6 | E | - | - | N | N | ns | + | - | - | P | E | ns |
| Atrazine | - | + | - | P | E | ns | + | + | E | P | P | ns | E | + | - | P | E | ns |
| Pyrimethanil | + | + | + | P | P | 1/6 | + | - | - | NA | E | ns | + | + | E | NA | P | 1/6 |
| Chlorpyrifos-methyl | + | + | + | P | P | 1/6 | + | B | - | NA | E | ns | + | + | E | NA | P | ns |

### 3.4 Gene expression measurement performance

Using a 2-fold mRNA threshold and the DIP classification based on batch calls, agreement with expected values was ≥80% for CYP1A2 and CYP3A4 and ∼78% for CYP2B6. Gene expression often clarified borderline/equivocal LC-MS/MS cases (e.g., rifampicin for CYP2B6), supporting its role as a cost-effective pre-screen and a supportive endpoint alongside enzymatic activity.

Across all chemicals and batches, gene expression demonstrated high concordance with expected classifications for CYP2B6 (13/16; 81.25%) and CYP3A4 (15/17; 88.2%), with CYP1A2 slightly below the ≥80% benchmark (14/19; 73.7%) (Tab. 7; Tab. S8). However, there is higher batch variability in the mRNA expression, as such it should only be used as a pre-screening approach.

**Tab. 7:** Gene expression classification of all chemicals. The gene expression upregulation fold-change results for three independent cell batches for treatment of 10 pharmaceutical proficiency, 6 augmentation and 4 additional chemicals compared against expected CYP activity from OECD draft TG and Jacobs et al, 2022. Green: agreement. CYP induction per batch indicated. (+: Positive; −: Negative; E: Equivocal; B: Borderline; P: Inducer; N: Non-inducer; #: Excluded from proficiency testing based on the validation ring trial results (JRC, 2014). Pre: Predicted classification; Res: Result from study; Classif: Classification Positive/ Negative / Equivocal. Significance assessed using one-way ANOVA with number of significant data points (n = 6). See Tab. S8. P ≤ 0.05; ** = P ≤ 0.01.

| Chemical | CYP1A2 |  |  |  |  | CYP2B6 |  |  |  |  | CYP3A4 |  |  |  |  |
| --- | --- | --- | --- | --- | --- | --- | --- | --- | --- | --- | --- | --- | --- | --- | --- |
|  | Batches |  |  | Classif. |  | Batches |  |  | Classif. |  | Batches |  |  | Classif. |  |
|  |  |  |  | Pre d. | Res . |  |  |  | Pre d. | Res . |  |  |  | Pre d. | Res . |
| Omeprazole | + | + | + | P | P | + | - | + | N# | P# | + | + | + | N | P |
| Carbamazepine | B | B | E | P | B | + | + | + | P | P | + | + | + | P | P |
| Phenytoin Na | B | E | B | P | B | + | + | + | P | P | + | + | + | P | P |
| Penicillin G | - | E | - | N | N | - | + | - | N | E | - | E | E | N | E |
| Sulfinpyrazone | B | + | + | P | P | + | + | + | P | P | + | + | + | P | P |
| Bosentan | + | + | - | P | P | + | B | - | P | E | + | + | + | P | P |
| Artemisinin | - | - | - | N | N | E | - | + | P | E | + | + | + | N | P |
| Rifampicin | + | - | + | P | P | + | + | + | P | P | + | + | + | P | P |
| Metoprolol | - | - | - | N | N | + | - | E | N | E | - | - | - | N | N |
| Sotalol | - | - | - | N | N | - | - | - | N | N | - | - | - | N | N |
| Tebuconazole | + | + | + | P | P | + | + | + | P | P | + | + | + | P | P |
| Benfuracarb | + | B | + | P | P | + | + | + | P | P | + | + | + | P | P |
| Chlorpyrifos | + | + | + | P | P | - | E | - | I | N | + | + | - | P | P |
| DEET | - | - | E | P | N | + | + | - | P | P | + | + | + | P | P |
| Fipronil | - | E | - | P | N | - | - | + | NA | E | + | + | + | P | P |
| Permethrin | E | E | + | N | E | E | E | - | N | E | - | - | B | NA | N |
| Prochloraz | + | + | + | NA | P | + | + | + | N | P | + | + | B | P | P |
| Atrazine | + | - | B | P | E | + | + | B | P | P | + | + | B | P | P |
| Pyrimethanil | + | + | + | P | P | + | E | + | NA | P | + | + | + | NA | P |
| Chlorpyrifos-methyl | E | - | - | P | N | E | E | - | NA | E | E | E | - | NA | E |

**Tab. 8:** Updated Data Interpretation Procedure (DIP) for the CYP induction HepaRG™ Test Guideline. Scenario on the classification of a chemical (in three independent experiments using three cell batches).

| Scenario | Experiment 1 | Experiment 2 | Experiment 3 | Conclusion |
| --- | --- | --- | --- | --- |
| 1 | positive | positive | positive | positive |
| 2 | positive | positive | borderline positive/<br>equivocal/ negative | positive |
| 3 | positive | borderline positive | borderline positive | positive |
| 4 | borderline positive | borderline positive | borderline positive | borderline positive |
| 5 | borderline positive | borderline positive | equivocal/negative | borderline positive |
| 6 | positive | borderline positive | equivocal/negative | equivocal |
| 7 | positive | equivocal/negative | equivocal/negative | equivocal |
| 8 | equivocal | equivocal | equivocal | equivocal |
| 9 | equivocal | equivocal | borderline positive/ negative | equivocal |
| 10 | negative | negative | borderline positive/<br>equivocal/ | negative |
| 11 | negative | negative | negative | negative |

**Tab. 9:** Between Laboratory Reproducibility for all proficiency chemicals. Comparison of the multi-classifier and two classifier DIP. Pharmaceuticals (n = 10) based on five laboratories (Bernasconi et al., 2019) and INRAE (Jacobs et al., 2026) and UKHSA. Augmentation chemicals (n = 6) based on two laboratories; UKHSA and best performing laboratory INRAE (less batch variability) in part 1 (Jacobs et al., 2026).

| Pharmaceuticals (five best performing laboratories) | Augmentation chemicals<br>(two best performing laboratories) |
| --- | --- |
| Multi-classifier DIP (positive; equivocal; borderline; negative) |  |
| CYP1A2 / 3A4 = 80%<br>CYP2B6 = 66% | CYP3A4 = 87.5%<br>CYP2B6 = 80%<br>CYP1A2 = 68.8% |
| Two classifier DIP (positive; negative) |  |
| CYP3A4 = 100%<br>CYP2B6 = 89%<br>CYP1A2 = 80% | CYP1A2 / 3A4 = 93.75%<br>CYP2B6 = 93.4% |

Pharmaceutical proficiency chemicals were more reproducible and robust using the enzymatic activity assessment, with an evident decrease in significance (Tab. 7; Fig. S3A-J). Notably, all additional chemicals aligned with predictions for CYP2B6 except prochloraz, which increased mRNA ∼10 x (Fig. 3A). CYP3A4 data were fully consistent between enzymatic activity and mRNA calls. For CYP1A2, discordances with DEET, fipronil, and chlorpyrifos-methyl, (no effect or equivocal induction to mRNA) despite predicted induction, were likely due to CYP1A2 exhibiting greater batch sensitivity in this test system, but also the weak/uncertain inducer properties of these chemicals (Jacobs et al., 2022) (Fig. 2A-F, 3A-D). The data indicate that mRNA measurements provide ≥80% agreement for CYP2B6/3A4 and can clarify borderline CYP activity calls and serve as a cost-efficient pre-screen.

**Fig. 2:**
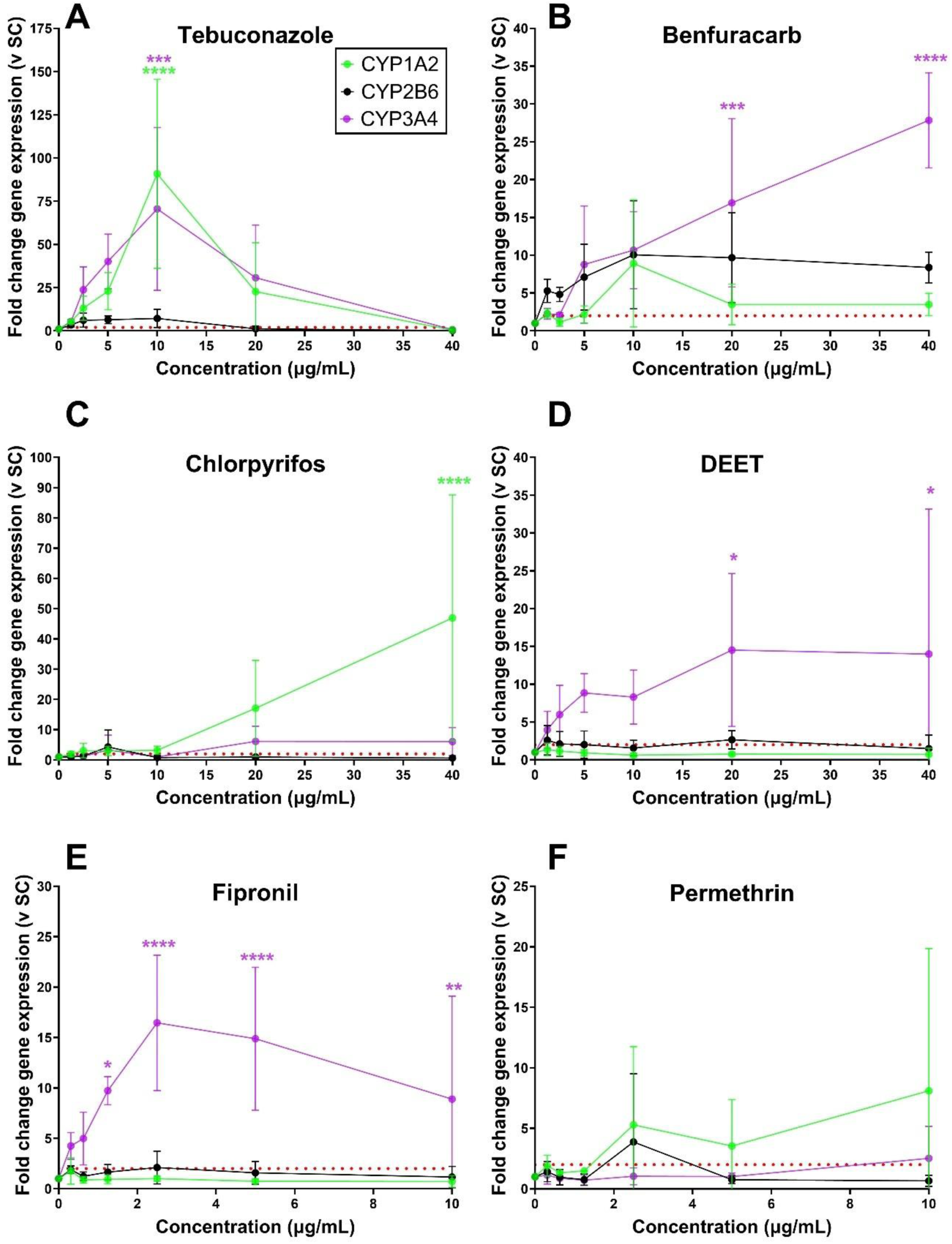
Gene expression fold induction concentration response for six augmentation chemicals tested. CYP1A2 (green), CYP2B6 (black) and CYP3A4 (purple) are shown. (A) Tebuconazole, (B) Benfuracarb, (C) Chlorpyrifos, (D) DEET, (E) Permethrin, (F) Fipronil. Three independent replicates (n = 3) (mean ± SD). Significance was assessed using one-way ANOVA * = P ≤ 0.05; ** = P ≤ 0.01; *** = P ≤ 0.001; **** = P ≤ 0.0001. GraphPad Prism Software v11.

**Fig. 3:**
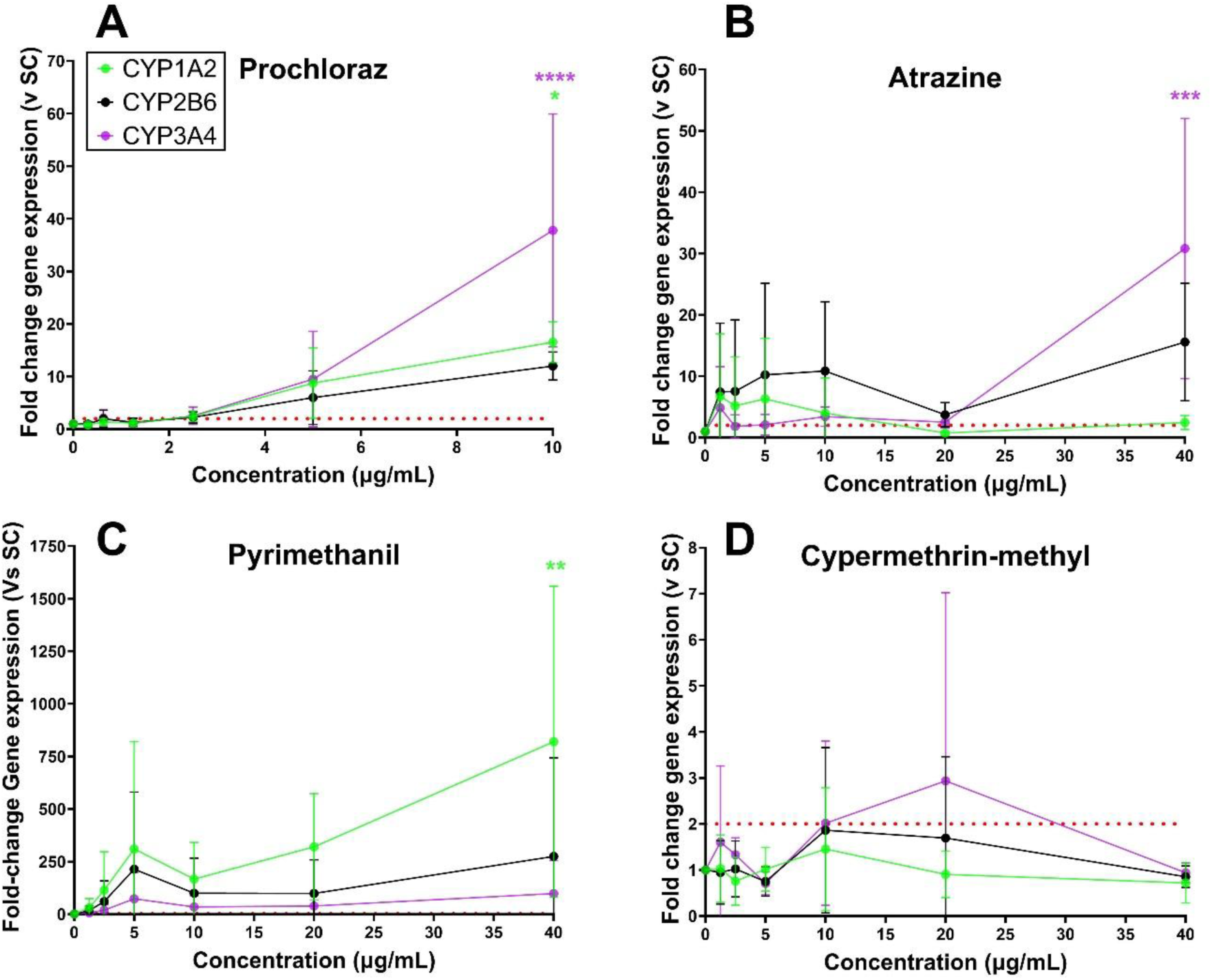
Gene expression fold induction concentration response for four additional chemicals tested. CYP1A2 (green), CYP2B6 (black) and CYP3A4 (purple) are shown. (A) Prochloraz, (B) Atrazine, (C) Pyrimethanil, (D) Chlorpyrifos-methyl. Three independent replicates (n = 3) (mean ± SD). * = P ≤ 0.05; ** = P ≤ 0.01; *** = P ≤ 0.001; **** = P ≤ 0.0001. GraphPad Prism Software v11.

## 4. Discussion and conclusion

### 4.1 Evaluation of Data Interpretation Procedures (DIPs)

The classification of CYP induction outcomes requires a DIP, and the data generated from the part 1 (Jacobs et al., 2026 submitted) and part 2 studies (this study) provides an opportunity to evaluate how different DIPs affect inducer calls. Two DIP systems were examined, the two classifier (JRC., 2014a; JRC TSAR protocol 2018; Bernasconi et al., 2019) DIP, and the updated multi-classifier DIP developed following the GOLIATH recommendations (Jacobs et al., 2026 submitted). In addition, appropriate statistical approaches contributed to decision robustness.

#### 4.1.1 Two classifier DIP

The original validation study (JRC., 2014a; Bernasconi et al., 2019) defines CYP induction as occurring when ≥2-fold activity is observed in two consecutive concentrations in at least one of three independent cell batches. This approach makes the test highly sensitive, because a positive outcome in only a single batch is sufficient to classify a chemical as an inducer. The 2019 DIP yielded 100% concordance for CYP1A2 and CYP3A4 and 100% for CYP2B6 (9/9) for pharmaceutical proficiency chemicals, confirming that the laboratory could successfully establish the validated method. However, the same DIP also contributed to false-positive outcomes in the wider GOLIATH dataset, particularly for CYP2B6, because a single-batch upward fluctuation could trigger an inducer classification. Thus, while the two classifier DIP supports assay transferability, as previously noted (JRC 2014b, c, Jacobs et al., 2026 submitted), it may overrepresent induction in cases where results are borderline or there is batch variability.

#### 4.1.2 Multi-classifier DIP: Positive, Borderline, Equivocal, Negative

To address these concerns, a more structured multi-classifier DIP was introduced as first described in Jacobs et al., part 1, and then further refined in discussion with European regulatory agencies, adding new outcome categories: positive, borderline, equivocal, and negative. The updated scheme applies more restrictive rules with respect to the classification of a test chemical as an *in vitro* CYP enzyme activity inducer. The four criteria outlined in the new DIP must be met to determine a positive, equivocal, borderline positive or negative classification.

Based upon this study alone, each of the chemicals evaluated resulted in enzymatic induction aligned with prior prediction (Bernasconi et al., 2019; Jacobs et al., 2022), where classifiers were either inducer (positive, borderline positive, equivocal) or no effect (negative). The introduction of borderline and equivocal inducers, for weaker hits, together with the recommendation of additional batch testing to increase data robustness for classification, improves the evidence base for when batch variability is an issue. These revisions support the updated DIP’s ability to reduce false positives by flagging uncertain or variable results rather than forcing overinterpretation of the results.

The between laboratory reproducibility (BLR) was evaluated and is shown in Tab.9. For the pharmaceutical proficiency chemicals, comparative concordance with the three JRC validation laboratories (Bernasconi et al., 2019) and INRAE (Jacobs et al., 2026) concordance was 80% for CYP1A2 and CYP3A4, with lower confidence for CYP2B6 across laboratories. Augmentation chemicals for two laboratories (INRAE and UKHSA) indicated concordance above 80% for CYP2B6 and CYP3A4, with lower confidence for CYP1A2. Overall, the studies conducted provide reliable enzymatic induction data regarding CYP1A2 and CYP3A4, but with less certainty than for CYP3A4, and this is therefore indicated in the respective TG445A (OECD., 2026).

As an encouraging note, it is expected that the variability between cell batches will reduce, as the manufacturers report that they have ascertained the likely cause of the variability during production, and that they can remediate this.

### 4.2 Gene expression endpoint role

In parallel with CYP activity measurements, exploratory CYP1A2, CYP2B6 and CYP3A4 gene-expression analysis was conducted only in the UKHSA laboratory. This was neither part of the originally validated CYP induction HepaRG^TM^ test method, nor was it subjected to formal inter-laboratory validation. As explicitly noted in methods (section 2.5), this is an altered unvalidated, investigational measurement and endpoint, rather than a validated component of the established test method and adopted TG.

Although not validated, the parallel gene-expression dataset may provide valuable mechanistic insight that can address certain limitations inherent in relying solely on CYP enzymatic activity. Enzyme activity integrates both *de novo* protein synthesis and protein stabilization as well as allostery, but it may not fully capture upstream receptor or other transcriptional activation events—especially for chemicals with low metabolic turnover, weak induction potencies, or concentration–response profiles affected by batch variability. By contrast, mRNA induction reflects direct nuclear receptor activation (AhR for CYP1A2; CAR for CYP2B6; PXR for CYP3A4), allowing earlier or more subtle biological responses to be detected. These complementary properties help reduce the risk of misclassifying borderline or weak inducers when enzyme activity (as determined by LC-MS/MS) alone is used as the decision driver.

The gene-expression results demonstrated ≥80% concordance with expected outcomes for CYP2B6 and CYP3A4, and 72.2% for CYP1A2, indicating reasonable mechanistic sensitivity. The data further showed that augmentation and additional chemicals generally behaved as predicted, although some discrepancies occurred (e.g., DEET, fipronil, chlorpyrifos -methyl). These findings suggest potential utility for gene-expression measures, as conducted here, as a useful support tool, especially when assay batches show technical variability in enzyme activity. Incorporating mRNA-based induction endpoints is consistent with current U.S. FDA drug–drug interaction (DDI) guidance (FDA, 2024), which recognizes CYP mRNA upregulation in human-relevant hepatic systems as an acceptable indicator of induction potential, provided it is interpreted within a weight-of-evidence framework. The HepaRG™ gene-expression data therefore aligns conceptually with international pharmacological regulatory expectations.

Overall, while gene-expression analysis cannot at this stage be used for definitive classification or OECD TG-aligned regulatory decisions, its application illustrates its potential added value in future method development. It may help strengthen IATAs by providing mechanistic confidence, improving sensitivity for weak inducers, and reducing uncertainty associated with enzyme-only measurements. However, for TG purposes, under the global regulatory OECD Test Guideline Mutual Acceptance of Data Agreement, further inter-laboratory validation using optimized and harmonized SOPs, will be required before it can be integrated formally into an OECD TG.

### 4.3. Limitations of the chemical augmentation and validation study

The validation study design addresses only measurement of CYP induction, however the method is also able to address CYP inhibition, which can be particularly important for drug-drug interactions and pharmaceutical efficacy. This was noted specifically for bosentan hydrate and chlorpyrifos (CYP2B6), artemisinin (CYP1A2) and omeprazole (CYP3A4), as indicated in Tab S5B. Bosentan hydrate with an absence of human *in vivo* data, was predicted to be a positive inducer in HepaRG cells (Bernasconi et al., 2019), yet it is evident there is a statistical significant (p ≤ 0.0001) decline in enzymatic activity, in agreement with GOLIATH laboratories (Jacobs et al 2026 submitted). Chlorpyrifos was the only defined inhibitor across the chemical set tested (Abass, Lämsä et al., 2012, 2013; D’Agostino, Zhang et al., 2015; Jacobs et al., 2022), where it was shown to be a strong inhibitor in the HepaRG cells (Tab. S5B, 6; Fig. 2C). Artemisinin moderately inhibits CYP1A2 at higher concentrations which also aligns with previously published data (Ericsson et al., 2014) (Tab. S5A). Omeprazole weakly inhibited CYP3A4 within two cell batches, which has also been reported for in drug-drug interaction studies (Shirasaka, Y et al., 2013). Future validation work for CYP inhibition would be required with additional known inhibitors used along with determination of IC_50_ values for a comprehensive targeted approach.

### 4. 4 Regulatory Applications of this test method

In addition to the generation of new reproducible augmented chemical applicability domain data, the WNT requested illustrations of applications of the test method, in integrated testing paradigms.

Suitable applications were identified in 2008 (OECD 2008, republished in 2014) and in relation to a call for the follow-up and validation of *in vitro* test methods for human metabolism, in particular to support endocrine active substance (EAS) hazard assessment, in order to facilitate the shift from *in vivo* to *in vitro* testing paradigms, by reducing the qualitative and quantitative deficiencies of *in vitro* EAS TGs (Jacobs et al 2013). This is viewed as particularly important for the testing of EAS, as several hormonally active chemicals, including some that occur naturally, are known to require bioactivation. The 2013 paper provided an illustration of how and where metabolism data, and in particular CYP induction data can be integrated into the OECD Conceptual Framework for the Testing and Assessment of Endocrine Disruptors (OECD 2018b), and a further update to this was provided in Jacobs et al 2022.The test method also has a fundamental role to play in most of the complex human health endpoints IATA, including for example the OECD Non-Genotoxic Carcinogen IATA (Jacobs et al 2020, Louekari and Jacobs 2024), where it is placed in Module B.

It has been placed similarly in relation to an IATA and integrated Adverse Outcome Pathway analysis for thyroid disruption (Vergauwen et al 2026 submitted), and for metabolic disruption (Jacobs/Ozcagli et al in prep).

Another key utility of the data will be for MetaPath. The US EPA’s MetaPath tool is a database of xenobiotic metabolism pathways (Kolanczyk et al., 2012). It is a knowledge-based expert system that can store and analyze summary information on pesticide metabolites found in several species (rodents, livestock, plants), residues of concern, and environmental degradates. The information is obtained from expert reviews of registrant-submitted studies as found in US, Canadian Data Evaluation Records (DERs) and European Draft Assessment Reports (DARs) and the published Assessment Reports, often according to the OECD Toxicokinetics TG 417 (OECD 2010). Efforts to expand MetaPath’s data population have been supported by many national pesticide regulatory bodies, and applicants submitting an application under Regulation (EC) No 1107/2009 are now requested by the European Food Safety Authority (EFSA) to provide data on metabolism in the area of residues and mammalian toxicology using the Metapath composer software. (https://www.efsa.europa.eu/en/applications/pesticides/tools, accessed 5 June 2026). With the eventual inclusion of human CYP induction data, it could become an excellent tool for simulating metabolism not only across species, but also in relation to human Phase II metabolism. With respect to PBPK tools commonly utilised by pharma, it will also provide useful data for SIMCYP (https://www.certara.com/software/simcyp-pbpk/ accessed 08/05/2026) and other *in silico* PBPK modelling initiatives. However an important caveat is that to date, the test method has not been developed to identify test chemical metabolites, but is a development that could be met in the future.

The recent FDA recommendations (FDA 2024) will also be well supported with the availability of a CYP induction TG.

Bringing the experimental work together, reported herein and in part 1, with the development and adoption of a rigorous DIP, and following peer review within the relevant OECD expert group, a revised draft TG has now been adopted by the WNT in April 2026. Together with Bernasconi et al., 2019, these part 1 (Jacobs et al., 2026 submitted) and part 2 papers provide the additional evidence basis required for the adoption.

## Disclaimer

The views and opinions expressed in this article are those of the author(s) and are not necessarily those of UK Health Security Agency or the Department of Health and Social Care.

## Declaration of generative AI in scientific writing

Microsoft M365 Copilot was used to assist with initial manuscript structuring and preparation of the plain-language summary. The authors have fully reviewed, revised and take full responsibility for the content.

## Acknowledgements

First and foremost, the critical validation work conducted by the JRC, led by Sandra Coecke and Camilla Bernasconi, is acknowledged.

The authors also gratefully acknowledge the preceding chemical augmentation laboratory work conducted under the auspices of the H2020 EU funded GOLIATH consortium, INRAE and Utrecht, learnings from which, contributed to the study design.

The active peer review of the OECD Expert Group for Toxicokinetics is gratefully acknowledged; and particularly in relation to the redrafting of the OECD TG 445A that was adopted in April 2026. The support also of Hugo Coppens and Christophe Chésne of Biopredic International, particularly in responding to some of the queries raised during the OECD review process is gratefully acknowledged.

## Funding

UK Government Department for Environment, Food and Rural Affairs and UK Health Security Agency Department of Toxicology.

## Notes

### Competing Interest Statement

The authors have declared no competing interest.

