## Supplementary information for "Chemical augmentation of the validated HepaRG^TM^ CYP induction test method Part 2: Additional laboratory study supported by mRNA analysis"

*Quartermain et al.:*

**Supplementary Figures and Tables**

Green: test chemical A (tested at six concentrations)

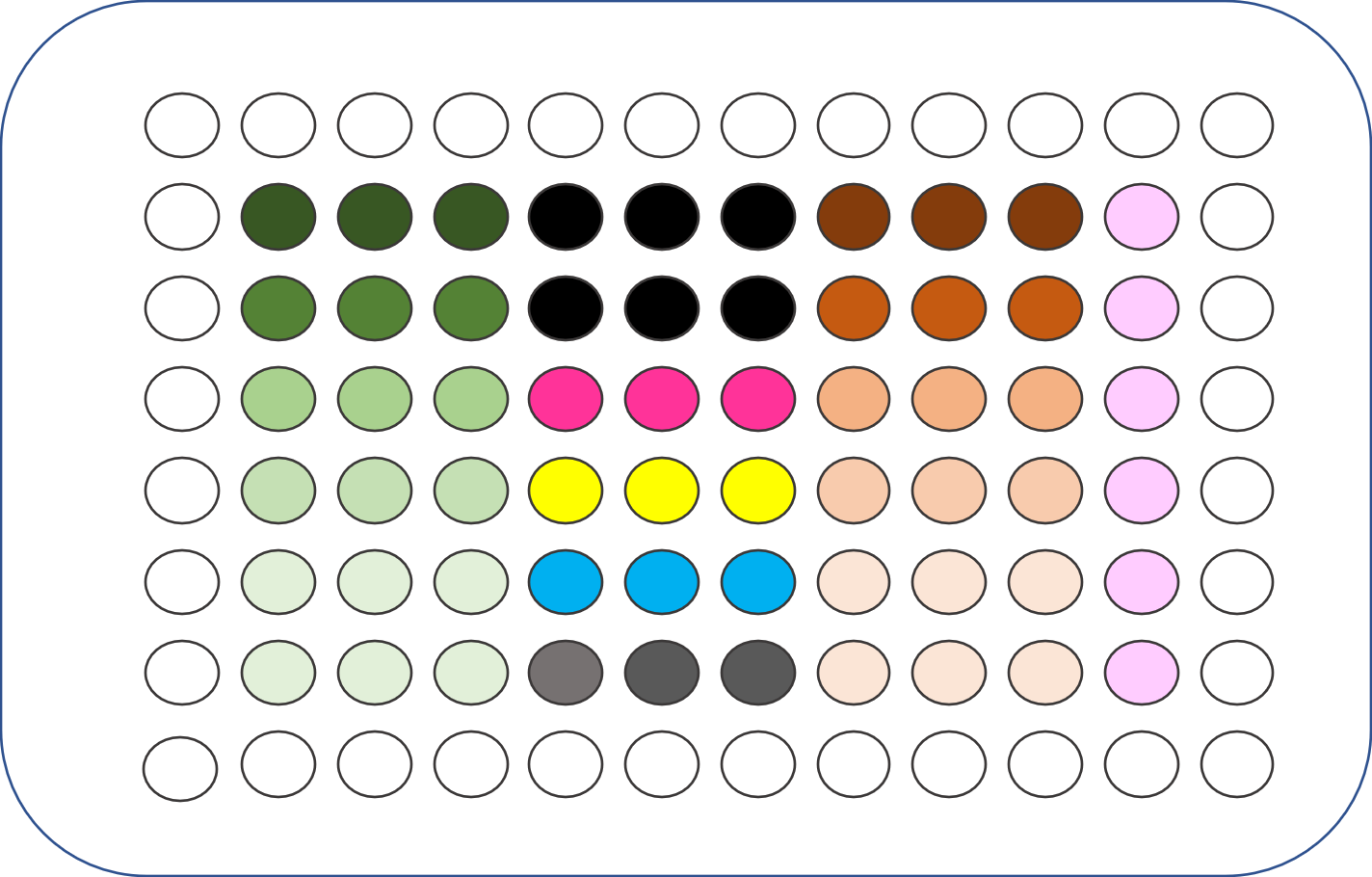
Orange: test chemical B (tested at six concentrations)

Black: solvent 0.1% DMSO test chemical A and B

Pink: reference chemical (positive control CYP2B6): phenobarbital 500 µM

Yellow: reference chemical (positive control CYP1A2):

β-naphthoflavone 25 µm

Blue: reference chemical (positive control CYP3A4): rifampicin 10 µM

Grey: solvent 0.1% DMSO control reference items

Pale pink: solvent free control

**Fig. S1: Plate layout for treatments with induction solutions.**

Shows test chemical A and B; 0.1% solvent control; reference chemicals Phenobarbital, β-naphthoflavone and Rifampicin; solvent free control. Three technical replicates are shown on each plate.

**Tab. S1: Calibration standards preparation in incubation medium**.

Calibration standards (CS) are prepared by fortifying the matrix (Incubation medium) with intermediate solutions of metabolites of the three metabolites 1’-Hydroxymidazolam (OH-MID), Hydroxybupropion (OH-BUP), Acetaminophen (ACETA) in acetonitrile.

| **Calibration standards (CS)** | | |
| --- | --- | --- |
| **N°** | **Concentration (pg/µL)** | |
| **CS1** | **OH-MID** | 1,400 |
|  | **OH-BUP** | 280 |
|  | **ACETA** | 280 |
| **CS2** | **OH-MID** | 700 |
|  | **OH-BUP** | 140 |
|  | **ACETA** | 140 |
| **CS3** | **OH-MID** | 350 |
|  | **OH-BUP** | 70 |
|  | **ACETA** | 70 |
| **CS4** | **OH-MID** | 175 |
|  | **OH-BUP** | 35 |
|  | **ACETA** | 35 |
| **CS5** | **OH-MID** | 87.5 |
|  | **OH-BUP** | 17.5 |
|  | **ACETA** | 17.5 |
| **CS6** | **OH-MID** | 43.75 |
|  | **OH-BUP** | 8.75 |
|  | **ACETA** | 8.75 |
| **CS7** | **OH-MID** | 21.875 |
|  | **OH-BUP** | 4.375 |
|  | **ACETA** | 4.375 |
| **CS8** | **OH-MID** | 10.9375 |
|  | **OH-BUP** | 2.1875 |
|  | **ACETA** | 2.1875 |
| **CS9** | **OH-MID** | 4.375 |
|  | **OH-BUP** | 0.875 |
|  | **ACETA** | 0.875 |

**Tab. S2.: Quality Control standards preparation in incubation medium**.

Quality Controls (QC) are independently prepared in the matrix from independent intermediate solutions of metabolites in acetonitrile.

| **QC preparation** | | |
| --- | --- | --- |
| **N°** | **Concentration (pg/µL)** | |
| **QC1-High** | **OH-MID** | 700 |
|  | **OH-BUP** | 140 |
|  | **ACETA** | 140 |
| **QC2-Mid** | **OH-MID** | 350 |
|  | **OH-BUP** | 70 |
|  | **ACETA** | 70 |
| **QC3-Low** | **OH-MID** | 21.875 |
|  | **OH-BUP** | 4.375 |
|  | **ACETA** | 4.375 |

**Tab. S3: Primer sequences used for gene expression analysis**.

TATA-binding protein (TBP) was used as the housekeeper gene (Housekeeper) to normalise using ΔΔCt method. Phase I genes for CYP1A2, CYP2B6 and CYP3A4 were designed in PrimerQuest, and their forward and reverse sequences are shown.

| Primer Sequences (5’-3’) | | |
| --- | --- | --- |
| Gene Name | **Forward primer sequence** | **Reverse primer sequence** |
| TBP | GAG CTG TGA TGT GAA GTT TCC | TCT GGG TTT GAT CAT TCT GTA G |
| CYP1A2 | CAG GAG CAC TAT CAG GAC TTT G | AAT GCT CCA GGT GAT GGC TGT G |
| CYP2B6 | CAC GCT CTC GCT CTT CTT T | CTC TCT GCA ACA TGA GGG TAT T |
| CYP3A4 | ACA GTC TTT CCA TTC CTC ATC C | GTA TCT TCG AGG CGA CTT TCT T |

**Tab. S4: Batch specific positive control fold induction as compared with solvent control for all tested chemicals**.

(A) β-naphthoflavone n-fold change induction for CYP1A2; (B) Phenobarbital n-fold induction for CYP2B6; (C) Rifampicin n-fold induction for CYP3A4. The HPR116 cell batch is indicated. Experimental runs must be at least, or greater than a 2-fold-change to be accepted. Based on two chemicals run per plate, including SD, Mean and RSD to indicate batch variation. As can be seen all values are above 2 fold, as required.

A)

| **Chemical** | **HPR116 #308** | **HPR116 #332** | **HPR116 #344** | **HPR116 #345** | **HPR116 #338** | **HPR116 #358** |
| --- | --- | --- | --- | --- | --- | --- |
| Prochloraz |  | 105.51 |  |  | 3.43 | 3.88 |
| Atrazine |  | 105.51 |  |  | 3.43 | 3.88 |
| Pyrimethanil |  | 25.04 |  |  | 5.11 | 3.35 |
| Chlorpyrifos-methyl |  | 25.04 |  |  | 5.11 | 3.35 |
| Tebuconazole |  | 12.87 | 4.17 | 3.71 |  |  |
| Benfuracarb |  | 12.87 | 4.17 | 3.71 |  |  |
| Chlorpyrifos |  | 2.30 | 4.29 | 19.13 |  |  |
| DEET |  | 2.30 | 4.29 | 19.13 |  |  |
| Permethrin |  | 8.90 | 34.44 | 12.88 |  |  |
| Fipronil |  | 8.90 | 34.44 | 12.88 |  |  |
| Phenytoin Na | 2.04 |  | 4.47 | 6.22 |  |  |
| Penicillin G Na | 2.04 |  | 4.47 | 6.22 |  |  |
| Rifampicin | 6.11 |  | 10.51 | 11.53 |  |  |
| Metoprolol | 6.11 |  | 10.51 | 11.53 |  |  |
| Carbamazepine | 2.02 |  | 3.39 | 3.39 |  |  |
| Omeprazole | 2.02 |  | 3.39 | 3.39 |  |  |
| Artemisinin | 2.35 |  | 3.43 | 4.49 |  |  |
| Sulfinpyrazone | 2.35 |  | 3.43 | 4.49 |  |  |
| Sotalol | 3.35 |  | 8.20 | 13.21 |  |  |
| Bosentan | 3.35 |  | 8.20 | 13.21 |  |  |
| STANDARD DEV | 1.54 | 38.02 | 9.86 | 5.34 | 0.84 | 0.27 |
| mean | 3.17 | 30.92 | 9.11 | 9.32 | 4.27 | 3.62 |
| RSD | 48.67 | 122.95 | 108.25 | 57.35 | 19.61 | 7.42 |

B)

| **Chemical** | **HPR116 #308** | **HPR116 #332** | **HPR116 #344** | **HPR116 #345** | **HPR116 #338** | **HPR116 #358** |
| --- | --- | --- | --- | --- | --- | --- |
| Prochloraz |  | 8.96 |  |  | 5.31 | 8.42 |
| Atrazine |  | 8.96 |  |  | 5.31 | 8.42 |
| Pyrimethanil |  | 58.66 |  |  | 4.58 | 6.92 |
| Chlorpyrifos-methyl |  | 58.66 |  |  | 4.58 | 6.92 |
| Tebuconazole |  | 7.53 | 9.54 | 7.29 |  |  |
| Benfuracarb |  | 7.53 | 9.54 | 7.29 |  |  |
| Chlorpyrifos |  | 6.81 | 3.63 | 13.28 |  |  |
| DEET |  | 6.81 | 3.63 | 13.28 |  |  |
| Permethrin |  | 4.13 | 17.62 | 4.63 |  |  |
| Fipronil |  | 4.13 | 17.62 | 4.63 |  |  |
| Phenytoin Na | 3.80 |  | 6.10 | 7.91 |  |  |
| Penicillin G Na | 3.80 |  | 6.10 | 7.91 |  |  |
| Rifampicin | 2.73 |  | 3.94 | 2.80 |  |  |
| Metoprolol | 2.73 |  | 3.94 | 2.80 |  |  |
| Carbamazepine | 3.26 |  | 3.55 | 6.89 |  |  |
| Omeprazole | 3.26 |  | 3.55 | 6.89 |  |  |
| Artemisinin | 2.81 |  | 4.40 | 5.74 |  |  |
| Sulfinpyrazone | 2.81 |  | 4.40 | 5.74 |  |  |
| Sotalol | 9.00 |  | 12.34 | 12.38 |  |  |
| Bosentan | 9.00 |  | 12.34 | 12.38 |  |  |
| STANDARD DEV | 2.37 | 1.46 | 4.80 | 3.38 | 0.37 | 0.75 |
| mean | 4.32 | 6.15 | 7.64 | 7.62 | 4.94 | 7.67 |
| RSD | 54.87 | 23.77 | 62.89 | 44.33 | 7.45 | 9.78 |

C)

| **Chemical** | **HPR116 #308** | **HPR116 #332** | **HPR116 #344** | **HPR116 #345** | **HPR116 #338** | **HPR116 #358** |
| --- | --- | --- | --- | --- | --- | --- |
| Prochloraz |  | 25.17 |  |  | 9.56 | 9.34 |
| Atrazine |  | 25.17 |  |  | 9.56 | 9.34 |
| Pyrimethanil |  | 30.33 |  |  | 15.94 | 2.75 |
| Chlorpyrifos-methyl |  | 30.33 |  |  | 15.94 | 2.75 |
| Tebuconazole |  | 81.46 | 36.52 | 26.49 |  |  |
| Benfuracarb |  | 81.46 | 36.52 | 26.49 |  |  |
| Chlorpyrifos |  | 21.12 | 10.80 | 18.61 |  |  |
| DEET |  | 21.12 | 10.80 | 18.61 |  |  |
| Permethrin |  | 12.21 | 28.08 | 5.54 |  |  |
| Fipronil |  | 12.21 | 28.08 | 5.54 |  |  |
| Phenytoin Na | 7.80 |  | 2.73 | 2.30 |  |  |
| Penicillin G Na | 7.80 |  | 2.73 | 2.30 |  |  |
| Rifampicin | 8.02 |  | 9.78 | 9.10 |  |  |
| Metoprolol | 8.02 |  | 9.78 | 9.10 |  |  |
| Carbamazepine | 6.22 |  | 4.72 | 14.70 |  |  |
| Omeprazole | 6.22 |  | 4.72 | 14.70 |  |  |
| Artemisinin | 2.40 |  | 9.46 | 12.61 |  |  |
| Sulfinpyrazone | 2.40 |  | 9.46 | 12.61 |  |  |
| Sotalol | 5.90 |  | 2.13 | 7.71 |  |  |
| Bosentan | 5.90 |  | 2.13 | 7.71 |  |  |
| STANDARD DEV | 2.01 | 30.76 | 11.73 | 7.28 | 3.19 | 3.29 |
| mean | 6.07 | 38.26 | 13.03 | 12.13 | 12.75 | 6.05 |
| RSD | 33.18 | 80.39 | 90.04 | 59.99 | 25.02 | 54.48 |

**Supplementary Data Table 5. Statistical CYP enzymatic induction data for all chemicals**.

**Tab. S5A: Statistical significance of cell batches (n = 3) and combined for CYP1A2 enzymatic induction, based on three replicates.** A one-way ANOVA compared against the solvent control corrected to Dunnett at p>0.05. Green-shading indicates significance. Red indicates significant decrease. ns = P < 0.05; * = P ≤ 0.05; ** = P ≤ 0.01; *** = P ≤ 0.001; **** = P ≤ 0.0001; **** = p<0.0001. P = inducer; N = non-inducer; N/A = insufficent prediction.Ome = Omeprazole; Carb = Carbamazepine; Phen Na = Phenytoin Sodium; Bos = Bosentan; Sot = Sotalol; Rif = Rifampicin; Met = Metoprolol; Art = Artemisinin; Sulf = Sulfinpyrazone; Teb = Tebuconazole; Benf = Benfuracarb; Chlor = Chlorpyrifos; DEET = N,N-diethyl-m-toluamide; Fip = Fipronil; Perm = Permethrin; Pro = Prochloraz; Atra = Atrazine; Pyrim = Pyrimethanil; Chlor-m = Chlorpyrifos-methyl

| **Combined cell batch analysis** | | | | | | | | | | | | | | | | | | | | |
| --- | --- | --- | --- | --- | --- | --- | --- | --- | --- | --- | --- | --- | --- | --- | --- | --- | --- | --- | --- | --- |
| **Result** | **P** | **P** | **P** | **N** | **P** | **P** | **N** | **P** | **E** | **E** | **P** | **P** | **P** | **P** | **P** | **E** | **P** | **E** | **P** | **P** |
| **Test conc** | **Ome** | **Carb** | **Phen Na** | **Pen G Na** | **Sulf** | **Bos** | **Art** | **Rif** | **Met** | **Sot** | **Teb** | **Benf** | **Chlor** | **DEET** | **Fip** | **Perm** | **Pro** | **Atra** | **Pyrim** | **Chlor-m** |
| 6 (lowest) | *** | *ns* | *ns* | *ns* | *** | *ns* | *ns* | **** | *ns* | *ns* | *ns* | *ns* | *ns* | ***** | *ns* | *ns* | *ns* | *ns* | *ns* | *ns* |
| 5 | **** | *ns* | *** | *ns* | *ns* | **** | *ns* | **** | *ns* | *ns* | **** | *ns* | *ns* | **** | *ns* | *ns* | *ns* | *ns* | *ns* | *ns* |
| 4 | ***** | *ns* | *** | *ns* | *** | *** | *ns* | **** | *ns* | *ns* | **** | *** | *ns* | **** | *ns* | *ns* | *ns* | *ns* | *ns* | *ns* |
| 3 | ***** | **** | ***** | *ns* | *** | *** | *ns* | *** | *ns* | *ns* | ***** | *** | *ns* | **** | *ns* | *ns* | ***** | *ns* | *** | *** |
| 2 | **** | **** | **** | *ns* | *ns* | *ns* | *** ↓* | **** | *ns* | *ns* | *** | *ns* | *** | *** | *** | *ns* | *ns* | *ns* | *ns* | *ns* |
| 1 (highest) | *** | *** | ***** | *ns* | *ns* | **** | *** ↓* | *** | *ns* | *ns* | *ns* | *** | *** | *ns* | *ns* | *ns* | *ns* | *ns* | *ns* | *ns* |
| **Independent batch analysis** | | | | | | | | | | | | | | | | | | | | |
| **Result** | **P** | **P** | **P** | **N** | **P** | **P** | **N** | **P** | **E** | **E** | **P** | **P** | **P** | **P** | **P** | **E** | **P** | **E** | **P** | **P** |
| **Test conc** | **Ome** | **Carb** | **Phen Na** | **Pen G Na** | **Sulf** | **Bos** | **Art** | **Rif** | **Met** | **Sot** | **Teb** | **Benf** | **Chlor** | **DEET** | **Fip** | **Perm** | **Pro** | **Atra** | **Pyrim** | **Chlor-m** |
| **Exp 1** | | | | | | | | | | | | | | | | | | | | |
| 6 (lowest) | *ns* | *ns* | *ns* | *ns* | *ns* | *ns* | *ns* | *ns* | *ns* | *ns* | *ns* | *ns* | *ns* | *ns* | *ns* | *ns* | *ns* | *ns* | *ns* | *ns* |
| 5 | *ns* | *ns* | *ns* | *ns* | *ns* | *ns* | *ns* | *ns* | *ns* | *ns* | **** | *ns* | *ns* | *** | *ns* | *ns* | *ns* | *ns* | *ns* | *ns* |
| 4 | *ns* | *ns* | *ns* | *ns* | *ns* | *ns* | *ns* | **** | *ns* | *ns* | **** | *ns* | *ns* | **** | *ns* | *ns* | **** | *ns* | *ns* | *ns* |
| 3 | *** | *ns* | *ns* | *ns* | *ns* | *ns* | *ns* | *ns* | *ns* | **** | **** | *ns* | *ns* | *ns* | *ns* | *ns* | *ns* | *ns* | ***** | *** |
| 2 | *** | *ns* | ****** | *ns* | *ns* | *ns* | *ns* | *ns* | *ns* | *ns* | *ns* | *ns* | *ns* | *ns* | *ns* | *ns* | *ns* | *ns* | **** | *ns* |
| 1 (highest) | ***** | *ns* | **** | *ns* | *ns* | *ns* | *ns* | *** | *ns* | ***** | *ns* | *ns* | *ns* | *ns* | *ns* | *ns* | *ns* | *ns* | *ns* | *ns* |
| **Exp 2** | | | | | | | | | | | | | | | | | | | | |
| 6 (lowest) | *ns* | *ns* | *ns* | *ns* | *ns* | *ns* | *ns* | *ns* | *ns* | *ns* | **** | *ns* | *ns* | *ns* | *ns* | *ns* | *ns* | *ns* | *ns* | *ns* |
| 5 | *ns* | *ns* | *ns* | *ns* | *ns* | *ns* | *ns* | *ns* | *ns* | *ns* | ****** | *ns* | ***** | *ns* | *ns* | *ns* | *ns* | *** | *ns* | *ns* |
| 4 | *** | *ns* | *ns* | *ns* | *** | *ns* | *ns* | *ns* | *ns* | *ns* | *ns* | *ns* | *ns* | ***** | *ns* | *ns* | *ns* | *** | *ns* | *ns* |
| 3 | ***** | *ns* | *ns* | *ns* | *ns* | *ns* | *ns* | *ns* | *ns* | *ns* | *ns* | *ns* | *ns* | *ns* | *ns* | *ns* | *** | *ns* | *ns* | *ns* |
| 2 | ****** | ***** | *ns* | *ns* | *ns* | *ns* | *ns* | *ns* | *ns* | *ns* | *ns* | ***** | **** | ***** | *ns* | *ns* | *ns* | *ns* | *ns* | *ns* |
| 1 (highest) | ****** | ***** | *** | *ns* | *** | *ns* | *ns* | *ns* | *ns* | *ns* | *ns* | ****** | ****** | *ns* | *ns* | *ns* | *ns* | *ns* | *ns* | *ns* |
| **Exp 3** | | | | | | | | | | | | | | | | | | | | |
| 6 (lowest) | *ns* | **** | *ns* | *ns* | *** | ****** | *ns* | *ns* | *ns* | *ns* | *ns* | *ns* | *ns* | *ns* | *ns* | *ns* | *ns* | *ns* | *ns* | *ns* |
| 5 | *ns* | ****** | *ns* | *ns* | *ns* | ****** | *ns* | *ns* | *ns* | *ns* | *** | *ns* | *ns* | *ns* | *ns* | *ns* | *ns* | *ns* | *ns* | *** |
| 4 | **** | *** | *ns* | *ns* | ***** | ****** | *ns* | *** | *ns* | *ns* | *ns* | *ns* | *ns* | *ns* | *ns* | *ns* | *ns* | *ns* | *ns* | ***** |
| 3 | ****** | ***** | *ns* | *ns* | **** | ***** | *ns* | *** | *ns* | *ns* | *ns* | *ns* | *ns* | *** | ***** | *** | *** | *ns* | *ns* | *ns* |
| 2 | ****** | ****** | **** | *ns* | **** | *ns* | *ns* | *ns* | *ns* | *ns* | *ns* | *** | *ns* | *ns* | *** | *ns* | *ns* | *ns* | *ns* | *ns* |
| 1 (highest) | ****** | ****** | **** | *ns* | ***** | *ns* | *ns* | *ns* | *ns* | *ns* | *ns* | **** | *ns* | *ns* | *ns* | *ns* | *ns* | *ns* | *ns* | *ns* |

**Tab. S5B: Statistical significance of cell batches (n = 3) and combined for CYP2B6 enzymatic induction, based on three replicates.** A one-way ANOVA compared against the solvent control corrected to Dunnett at p>0.05. Green-shading indicates significance. Red indicates significant decrease. ns = P <0.05; * = P ≤ 0.05; ** = P ≤ 0.01; *** = P ≤ 0.001; **** = P ≤ 0.0001. P = inducer; N = non-inducer; N/A = insufficent prediction. Ome = Omeprazole; Carb = Carbamazepine; Phen Na = Phenytoin Sodium; Bos = Bosentan; Sot = Sotalol; Rif = Rifampicin; Met = Metoprolol; Art = Artemisinin; Sulf = Sulfinpyrazone; Teb = Tebuconazole; Benf = Benfuracarb; Chlor = Chlorpyrifos; DEET = N,N-diethyl-m-toluamide; Fip = Fipronil; Perm = Permethrin; Pro = Prochloraz; Atra = Atrazine; Pyrim = Pyrimethanil; Chlor-m = Chlorpyrifos-methyl

| **Combined cell batch analysis** | | | | | | | | | | | | | | | | | | | | |
| --- | --- | --- | --- | --- | --- | --- | --- | --- | --- | --- | --- | --- | --- | --- | --- | --- | --- | --- | --- | --- |
| **Result** | **E** | **P** | **P** | **N** | **P** | **E** | **P** | **E** | **N** | **N** | **P** | **P** | **N** | **P** | **E** | **E** | **N** | **P** | **E** | **E** |
| **Test conc** | **Ome** | **Carb** | **Phen Na** | **Pen G Na** | **Sulf** | **Bos** | **Art** | **Rif** | **Met** | **Sot** | **Teb** | **Benf** | **Chlor** | **DEET** | **Fip** | **Perm** | **Pro** | **Atra** | **Pyrim** | **Chlor-m** |
| 6 (lowest) | *ns* | *ns* | *** | *ns* | **** | *ns* | *ns* | *ns* | *ns* | *ns* | *ns* | *** | *ns* | ***** | *ns* | *ns* | *ns* | *ns* | *ns* | *ns* |
| 5 | *ns* | *ns* | **** | *ns* | *ns* | *ns* | *ns* | *ns* | *ns* | *ns* | *** | *** | ** ↓* | ***** | *ns* | *ns* | *ns* | *yes (p<0.1)* | *ns* | *ns* |
| 4 | *ns* | *ns* | ***** | *ns* | **** | *ns* | *ns* | *yes (p<0.1)* | *ns* | *ns* | **** | ***** | ***** ↓* | **** | *ns* | *ns* | *ns* | *ns* | *ns* | *ns* |
| 3 | *ns* | *ns* | ****** | *ns* | **** | *ns* | *ns* | *ns* | *ns* | *ns* | **** | **** | ***** ↓* | **** | *ns* | *ns* | *ns* | *ns* | *ns* | *ns* |
| 2 | *ns* | *** | **** | *ns* | *** | ***** ↓* | ****** | *ns* | *ns* | *ns* | **** | *** | ***** ↓* | *ns* | *ns* | *ns* | *ns* | *ns* | *ns* | *ns* |
| 1 (highest) | *ns* | *** | **** | *ns* | ***** | ***** ↓* | ****** | *ns* | *ns* | *ns* | *** | *** | ***** ↓* | *** | *ns* | *ns* | *ns* | *ns* | *ns* | *ns* |
| **Independent batch analysis** | | | | | | | | | | | | | | | | | | | | |
| **Result** | **E** | **P** | **P** | **N** | **P** | **E** | **P** | **E** | **N** | **N** | **P** | **P** | **N** | **P** | **E** | **E** | **N** | **P** | **E** | **E** |
| **Test conc** | **Ome** | **Carb** | **Phen Na** | **Pen G Na** | **Sulf** | **Bos** | **Art** | **Rif** | **Met** | **Sot** | **Teb** | **Benf** | **Chlor** | **DEET** | **Fip** | **Perm** | **Pro** | **Atra** | **Pyrim** | **Chlor-m** |
| **Exp 1** | | | | | | | | | | | | | | | | | | | | |
| 6 (lowest) | *ns* | *ns* | *ns* | *ns* | *ns* | *ns* | *ns* | *ns* | *ns* | *ns* | *ns* | *ns* | *ns* | *** | *ns* | *ns* | ****** | *ns* | *ns* | *ns* |
| 5 | *ns* | *ns* | **** | *ns* | ***** | *ns* | *ns* | *ns* | *ns* | *ns* | *ns* | *ns* | ** ↓* | ***** | *ns* | *ns* | *ns* | *ns* | ****** | **** |
| 4 | *ns* | *ns* | *** | *ns* | *ns* | *ns* | *ns* | *ns* | *ns* | *ns* | *ns* | *ns* | *** ↓* | *** | *ns* | *ns* | *ns* | *ns* | ***** | **** |
| 3 | *ns* | *ns* | ****** | *ns* | *ns* | *ns* | *ns* | *ns* | *ns* | **** | *ns* | *ns* | **** ↓* | *ns* | *ns* | *ns* | *ns* | *ns* | *ns* | ***** |
| 2 | *ns* | *ns* | ****** | *ns* | *ns* | ** ↓* | *ns* | *ns* | *ns* | *ns* | *ns* | *ns* | **** ↓* | *ns* | *ns* | *ns* | *ns* | *ns* | *ns* | *ns* |
| 1 (highest) | *ns* | *ns* | **** | *ns* | *ns* | *** ↓* | *ns* | *ns* | *ns* | *ns* | *ns* | *ns* | **** ↓* | *ns* | *ns* | *ns* | *ns* | *ns* | *ns* | *ns* |
| **Exp 2** | | | | | | | | | | | | | | | | | | | | |
| 6 (lowest) | *ns* | *ns* | *ns* | *ns* | *ns* | *** ↓* | *** | *ns* | *ns* | *ns* | **** | *ns* | *ns* | ****** | *ns* | *ns* | *ns* | *ns* | *ns* | *ns* |
| 5 | *ns* | *ns* | *ns* | *ns* | *ns* | *ns* | *ns* | *ns* | *ns* | *ns* | ****** | **** | *ns* | ***** | *ns* | *ns* | *ns* | *ns* | *ns* | *ns* |
| 4 | *ns* | *ns* | *ns* | *ns* | *ns* | *ns* | *ns* | *ns* | *ns* | *ns* | **** | *** | **** ↓* | ****** | *ns* | *ns* | *ns* | *yes (p<0.1)* | *ns* | *ns* |
| 3 | *ns* | *ns* | *** | *ns* | *ns* | *ns* | *ns* | *ns* | *ns* | *ns* | **** | *ns* | ***** ↓* | *ns* | *ns* | *ns* | *ns* | *ns* | *ns* | *ns* |
| 2 | *ns* | *ns* | *** | *ns* | *ns* | ** ↓* | *ns* | *ns* | *ns* | *ns* | *ns* | **** | ***** ↓* | ***** | *ns* | *ns* | *ns* | *ns* | *ns* | *ns* |
| 1 (highest) | *ns* | *ns* | *ns* | *ns* | *ns* | ** ↓* | *ns* | *ns* | *ns* | *ns* | *ns* | ****** | ***** ↓* | *ns* | *ns* | *ns* | *ns* | *ns* | *ns* | *ns* |
| **Exp 3** | | | | | | | | | | | | | | | | | | | | |
| 6 (lowest) | *ns* | ***** | *ns* | *ns* | *ns* | ***** ↓* | ****** | *ns* | *ns* | *ns* | *ns* | *ns* | *ns* | **** | *ns* | *ns* | *ns* | *ns* | *ns* | *ns* |
| 5 | *ns* | ****** | *ns* | *ns* | *ns* | *** ↓* | *ns* | *ns* | *ns* | *ns* | ***** | **** | ** ↓* | *** | *ns* | *ns* | *ns* | *ns* | *ns* | *ns* |
| 4 | **** | ****** | *** | *ns* | *ns* | ** ↓* | *ns* | *ns* | *ns* | *ns* | *** | *ns* | **** ↓* | *ns* | *ns* | *ns* | *ns* | *ns* | *ns* | *ns* |
| 3 | ****** | ****** | **** | *ns* | *ns* | *ns* | *ns* | *ns* | *ns* | *ns* | **** | *ns* | ***** ↓* | *ns* | *ns* | *** | *ns* | *ns* | *ns* | *ns* |
| 2 | ****** | ****** | **** | *ns* | *ns* | *ns* | *ns* | *ns* | *ns* | *ns* | *ns* | *ns* | **** ↓* | *ns* | *ns* | *ns* | *ns* | *ns* | *ns* | *ns* |
| 1 (highest) | *** | ****** | *ns* | *ns* | *ns* | ** ↓* | *ns* | *ns* | *ns* | *ns* | *ns* | *ns* | ***** ↓* | *ns* | *ns* | *ns* | *ns* | *ns* | *ns* | *ns* |

**Tab. S5C: Statistical significance of cell batches (n = 3) and combined for CYP3A4 enzymatic induction, based on three replicates**. A one-way ANOVA compared against the solvent control corrected to Dunnett at p>0.05. Green-shading indicates significance. Red indicates significant decrease. ns = P > 0.05; * = P ≤ 0.05; ** = P ≤ 0.01; *** = P ≤ 0.001; **** = P ≤ 0.0001. P = inducer; N = non-inducer; N/A = insufficent prediction. Ome = Omeprazole; Carb = Carbamazepine; Phen Na = Phenytoin Sodium; Bos = Bosentan; Sot = Sotalol; Rif = Rifampicin; Met = Metoprolol; Art = Artemisinin; Sulf = Sulfinpyrazone; Teb = Tebuconazole; Benf = Benfuracarb; Chlor = Chlorpyrifos; DEET = N,N-diethyl-m-toluamide; Fip = Fipronil; Perm = Permethrin; Pro = Prochloraz; Atra = Atrazine; Pyrim = Pyrimethanil; Chlor-m = Chlorpyrifos-m

| **Combined cell batch analysis** | | | | | | | | | | | | | | | | | | | | |
| --- | --- | --- | --- | --- | --- | --- | --- | --- | --- | --- | --- | --- | --- | --- | --- | --- | --- | --- | --- | --- |
| **Result** | **N** | **P** | **P** | **N** | **P** | **P** | **N** | **P** | **N** | **N** | **P** | **P** | **E** | **P** | **P** | **N** | **E** | **E** | **P** | **P** |
| **Test conc** | **Ome** | **Carb** | **Phen Na** | **Pen G Na** | **Sulf** | **Bos** | **Art** | **Rif** | **Met** | **Sot** | **Teb** | **Benf** | **Chlor** | **DEET** | **Fip** | **Perm** | **Pro** | **Atra** | **Pyrim** | **Chlor-m** |
| 6 (lowest) | *ns* | *ns* | *ns* | *ns* | ****** | *ns* | *ns* | ***** | *ns* | *ns* | *** | *ns* | *ns* | *** | *ns* | *ns* | *ns* | *ns* | *ns* | *ns* |
| 5 | *ns* | *ns* | *ns* | *ns* | *ns* | ***** | *ns* | **** | *ns* | *ns* | **** | *ns* | *** | **** | *ns* | *ns* | *ns* | *ns* | ****** | *ns* |
| 4 | *ns* | *** | **** | *ns* | ****** | *** | *ns* | **** | *ns* | *ns* | **** | **** | *** | **** | *** | *ns* | *ns* | *ns* | *ns* | *ns* |
| 3 | *ns* | ***** | **** | *ns* | ***** | **** | *ns* | ***** | *ns* | *ns* | ***** | **** | *ns* | **** | *ns* | *ns* | *ns* | *ns* | *ns* | *ns* |
| 2 | *ns* | *** | *** | *ns* | **** | *** | *ns* | ***** | *ns* | *ns* | **** | *** | *ns* | **** | **** | *ns* | *ns* | *ns* | *ns* | *ns* |
| 1 (highest) | *ns* | *** | **** | *ns* | ***** | *ns* | *ns* | **** | *ns* | *ns* | *ns* | *** | *** | ****** | *ns* | *ns* | *ns* | *ns* | *ns* | *ns* |
| **Independent batch analysis** | | | | | | | | | | | | | | | | | | | | |
| **Result** | **N** | **P** | **P** | **N** | **P** | **P** | **N** | **P** | **N** | **N** | **P** | **P** | **E** | **P** | **P** | **N** | **E** | **E** | **P** | **P** |
| **Test conc** | **Ome** | **Carb** | **Phen Na** | **Pen G Na** | **Sulf** | **Bos** | **Art** | **Rif** | **Met** | **Sot** | **Teb** | **Benf** | **Chlor** | **DEET** | **Fip** | **Perm** | **Pro** | **Atra** | **Pyrim** | **Chlor-m** |
| **Exp 1** | | | | | | | | | | | | | | | | | | | | |
| 6 (lowest) | *** ↓* | *ns* | *ns* | *ns* | *ns* | *ns* | *ns* | *ns* | *ns* | *ns* | *ns* | *ns* | *ns* | *ns* | *ns* | *ns* | *ns* | *ns* | *ns* | *ns* |
| 5 | *ns* | *ns* | *ns* | *ns* | **** | **** | *ns* | *ns* | *ns* | ***** | *ns* | *ns* | *ns* | ***** | *ns* | *ns* | **** | *ns* | *ns* | *ns* |
| 4 | *ns* | *ns* | *ns* | *ns* | *** | *ns* | *ns* | *ns* | *ns* | *ns* | *ns* | *ns* | *ns* | ****** | *ns* | *ns* | *ns* | *ns* | *ns* | *ns* |
| 3 | *ns* | *ns* | *ns* | *ns* | **** | ***** | *** | **** | *ns* | ***** | *** | *ns* | *ns* | ***** | *** | *ns* | *** | *ns* | *ns* | *ns* |
| 2 | *ns* | *ns* | **** | *ns* | **** | *ns* | *ns* | ***** | *ns* | *ns* | *ns* | *ns* | *ns* | *** | *ns* | *ns* | *** | *ns* | *ns* | *ns* |
| 1 (highest) | *ns* | *ns* | ****** | *ns* | ****** | *ns* | *** | ****** | *ns* | **** | *ns* | *ns* | *ns* | *ns* | *ns* | *ns* | *ns* | *ns* | *ns* | *ns* |
| **Exp 2** | | | | | | | | | | | | | | | | | | | | |
| 6 (lowest) | *ns* | *ns* | *ns* | *ns* | *ns* | *ns* | *ns* | *ns* | *ns* | *ns* | ****** | *ns* | *ns* | *ns* | *ns* | *ns* | *ns* | *ns* | *ns* | *ns* |
| 5 | *ns* | *ns* | *ns* | *ns* | *ns* | *** | *ns* | *ns* | *ns* | *ns* | ****** | *ns* | *ns* | *ns* | *ns* | *ns* | *ns* | *ns* | *ns* | *ns* |
| 4 | *ns* | *ns* | *ns* | *ns* | *ns* | **** | *ns* | ***** | *ns* | *ns* | ***** | *ns* | *ns* | ***** | *ns* | *ns* | *ns* | *ns* | *ns* | **** |
| 3 | *ns* | *ns* | *ns* | *ns* | *ns* | *** | *ns* | *** | *ns* | *ns* | *ns* | *ns* | *ns* | *ns* | *ns* | *ns* | *ns* | *ns* | *ns* | *ns* |
| 2 | *ns* | *ns* | *ns* | *ns* | *ns* | *ns* | *ns* | *ns* | *ns* | *ns* | *ns* | ****** | **** | ****** | *ns* | *ns* | *ns* | *ns* | *ns* | *ns* |
| 1 (highest) | *** ↓* | *** | *ns* | *ns* | *ns* | *ns* | *ns* | *ns* | *ns* | *ns* | *ns* | ****** | **** | *ns* | *ns* | *ns* | *ns* | *ns* | *ns* | *ns* |
| **Exp 3** | | | | | | | | | | | | | | | | | | | | |
| 6 (lowest) | *ns* | *ns* | *ns* | *ns* | *ns* | ****** | *ns* | *ns* | *ns* | *ns* | ***** | *ns* | *ns* | *ns* | *ns* | *ns* | *ns* | *ns* | *ns* | *ns* |
| 5 | *ns* | **** | *ns* | *ns* | *ns* | ****** | *ns* | *** | *ns* | *ns* | ****** | *ns* | *ns* | *ns* | *ns* | *ns* | *ns* | *ns* | *ns* | *ns* |
| 4 | *ns* | *ns* | *ns* | *ns* | *** | ****** | *ns* | ****** | *ns* | *ns* | **** | *ns* | *ns* | *** | *ns* | *ns* | *ns* | *ns* | *ns* | *ns* |
| 3 | *ns* | **** | *ns* | *ns* | ****** | ****** | *ns* | **** | *ns* | *ns* | *ns* | *ns* | *ns* | ***** | ****** | *ns* | *ns* | *ns* | *** | *ns* |
| 2 | *ns* | ****** | *ns* | *ns* | ****** | **** | *ns* | **** | *ns* | *ns* | *ns* | *** | *ns* | *ns* | *ns* | *ns* | *ns* | *ns* | *ns* | *ns* |
| 1 (highest) | *ns* | ****** | **** | *ns* | **** | *ns* | *ns* | *** | *ns* | *ns* | *ns* | ****** | *ns* | *ns* | *ns* | *ns* | *ns* | *ns* | *ns* | *ns* |

**Tab. S6: CYP enzymatic induction data for six proficiency augmentation chemicals against solvent control**.

CYP1A2, CYP2B6 and CYP3A4 are shown, where n = 3 cell batches mean ± SD.

**CYP1A2**

| **Tebuconazole** | | **Benfurcarb** | | **Chlorpyrifos** | | **DEET** | | **Permethrin** | | **Fipronil** | |
| --- | --- | --- | --- | --- | --- | --- | --- | --- | --- | --- | --- |
| µg/ml | n-fold induction (±SD) | µg/ml | n-fold induction (±SD) | µg/ml | n-fold induction (±SD) | µg/ml | n-fold induction (±SD) | µg/ml | n-fold induction (±SD) | µg/ml | n-fold induction (±SD) |
| 40 | 1.663 ± 1.022 | 40 | 7.833 ± 6.333 | 40 | 4.730 ± 3.238 | 40 | 1.162 ± 0.184 | 10 | 1.396 ± 0.567 | 10 | 1.794 ± 1.190 |
| 20 | 2.297 ± 1.227 | 20 | 5.421 ± 3.120 | 20 | 3.090 ± 1.790 | 20 | 2.957 ± 1.217 | 5 | 1.452 ± 0.884 | 5 | 3.451 ± 1.324 |
| 10 | 4.620 ± 1.800 | 10 | 2.497 ± 1.328 | 10 | 1.738 ± 0.744 | 10 | 2.758 ± 0.884 | 2.5 | 1.902 ± 1.000 | 2.5 | 3.704 ± 2.959 |
| 5 | 4.774 ± 1.968 | 5 | 2.537 ± 1.211 | 5 | 2.069 ± 1.051 | 5 | 3.596 ± 0.997 | 1.25 | 1.239 ± 0.464 | 1.25 | 2.870 ± 2.047 |
| 2.5 | 6.750 ± 1.449 | 2.5 | 2.553 ± 1.758 | 2.5 | 3.364 ± 2.429 | 2.5 | 2.796 ± 0.710 | 0.63 | 1.169 ± 0.584 | 0.63 | 2.084 ± 1.542 |
| 1.25 | 3.593 ± 1.022 | 1.25 | 1.759 ± 0.868 | 1.25 | 2.296 ± 1.135 | 1.25 | 1.681 ± 0.269 | 0.31 | 1.487 ± 0.449 | 0.31 | 2.083 ± 1.776 |

**CYP2B6**

| **Tebuconazole** | | **Benfurcarb** | | **Chlorpyrifos** | | **DEET** | | **Permethrin** | | **Fipronil** | |
| --- | --- | --- | --- | --- | --- | --- | --- | --- | --- | --- | --- |
| µg/ml | n-fold induction (±SD) | µg/ml | n-fold induction (±SD) | µg/ml | n-fold induction (±SD) | µg/ml | n-fold induction (±SD) | µg/ml | n-fold induction (±SD) | µg/ml | n-fold induction (±SD) |
| 40 | 3.238 ± 2.084 | 40 | 6.139 ± 3.834 | 40 | 0.271 ± 0.060 | 40 | 1.956 ± 0.419 | 10 | 1.349 ± 0.254 | 10 | 1.797 ± 1.224 |
| 20 | 3.912 ± 1.796 | 20 | 5.842 ± 2.231 | 20 | 0.270 ± 0.066 | 20 | 5.660 ± 4.030 | 5 | 1.266 ± 0.203 | 5 | 1.773 ± 0.176 |
| 10 | 6.423 ± 2.468 | 10 | 3.856 ± 1.375 | 10 | 0.193 ± 0.060 | 10 | 5.090 ± 1.018 | 2.5 | 1.616 ± 0.803 | 2.5 | 1.298 ± 0.887 |
| 5 | 5.912 ± 1.878 | 5 | 4.918 ± 1.799 | 5 | 0.308 ± 0.100 | 5 | 8.648 ± 3.740 | 1.25 | 1.009 ± 0.250 | 1.25 | 1.011 ± 0.589 |
| 2.5 | 7.727 ± 3.822 | 2.5 | 5.740 ± 3.500 | 2.5 | 0.629± 0.187 | 2.5 | 9.206 ± 1.443 | 0.63 | 0.830 ± 0.213 | 0.63 | 1.088 ± 0.649 |
| 1.25 | 4.474 ± 3.639 | 1.25 | 4.114 ± 2.069 | 1.25 | 0.713 ± 0.108 | 1.25 | 9.992 ± 3.816 | 0.31 | 1.318 ± 0.843 | 0.31 | 1.527 ± 1.400 |

**CYP3A4**

| **Tebuconazole** | | **Benfurcarb** | | **Chlorpyrifos** | | **DEET** | | **Permethrin** | | **Fipronil** | |
| --- | --- | --- | --- | --- | --- | --- | --- | --- | --- | --- | --- |
| µg/ml | n-fold induction (±SD) | µg/ml | n-fold induction (±SD) | µg/ml | n-fold induction (±SD) | µg/ml | n-fold induction (±SD) | µg/ml | n-fold induction (±SD) | µg/ml | n-fold induction (±SD) |
| 40 | 1.589 ± 1.072 | 40 | 16.344 ± 11.306 | 40 | 1.823 ± 0.735 | 40 | 3.444 ± 0.020 | 10 | 1.134 ± 0.598 | 10 | 2.66 ± 1.785 |
| 20 | 3.269 ± 0.888 | 20 | 10.730 ± 5.561 | 20 | 1.861 ± 0.784 | 20 | 7.144 ± 2.686 | 5 | 0.787 ± 0.256 | 5 | 7.973 ± 3.473 |
| 10 | 5.119 ± 1.047 | 10 | 4.523 ± 2.056 | 10 | 0.807 ± 0.203 | 10 | 6.918 ± 2.585 | 2.5 | 0.893 ± 0.358 | 2.5 | 15.368 ± 14.730 |
| 5 | 7.216 ± 2.351 | 5 | 4.026 ± 1.804 | 5 | 0.561 ± 0.073 | 5 | 8.926 ± 3.068 | 1.25 | 1.453 ± 0.672 | 1.25 | 7.863 ± 6.111 |
| 2.5 | 9.277 ± 4.602 | 2.5 | 3.370 ± 2.236 | 2.5 | 0.710 ± 0.170 | 2.5 | 6.119 ± 1.993 | 0.63 | 0.926 ± 0.257 | 0.63 | 3.839 ± 2.926 |
| 1.25 | 9.110 ± 6.468 | 1.25 | 1.118 ± 0.565 | 1.25 | 1.008 ± 0.296 | 1.25 | 1.403 ± 0.347 | 0.31 | 1.008 ± 0.122 | 0.31 | 1.723 ± 1.322 |

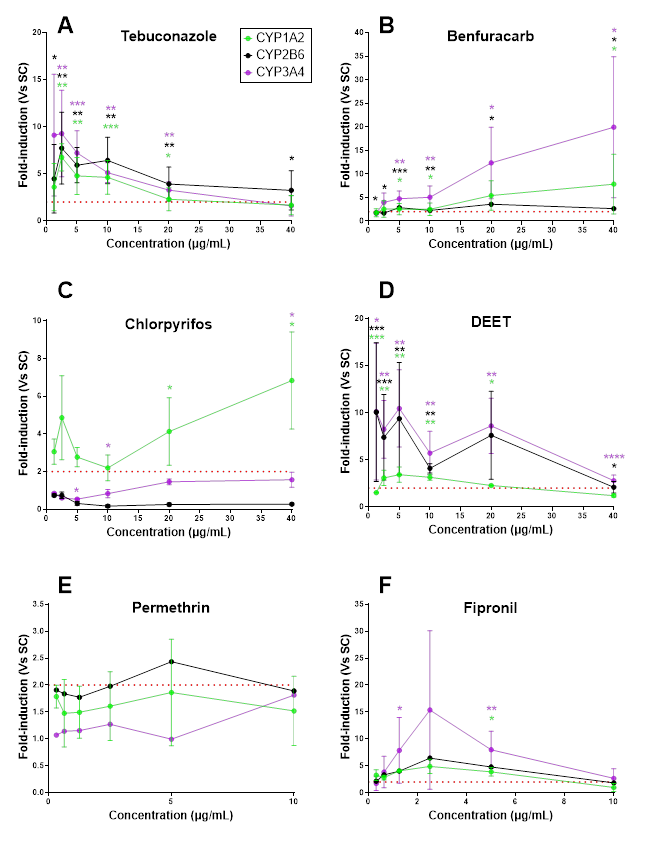

**Fig. S2A-F: Concentration response for augmentation chemicals tested.** CYP1A2 (green), CYP2B6 (black) and CYP3A4 (purple) are shown. (A) Tebuconazole; (B) Benfuracarb; (C) Chlorpyrifos; (D) DEET; (E) Permethrin; (F) Fipronil. Three independent replicates (n = 3) (mean ± SD). * = P ≤ 0.05; ** = P ≤ 0.01; *** = P ≤ 0.001; **** = P ≤ 0.0001. GraphPad Prism Software v11.

**Tab. S7: CYP enzymatic induction data for the four additional chemicals against solvent control**.

CYP1A2, CYP2B6 and CYP3A4 are shown, where n = 3 cell batches mean ± SD.

**CYP1A2**

| **Prochloraz** | | **Atrazine** | | **Pyrimethanil** | | **Chlorpyrifos-methyl** | |
| --- | --- | --- | --- | --- | --- | --- | --- |
| µg/ml | n-fold induction (±SD) | µg/ml | n-fold induction (±SD) | µg/ml | n-fold induction (±SD) | µg/ml | n-fold induction (±SD) |
| 10 | 3.554 ± 1.960 | 40 | 1.818 ± 2.904 | 40 | 31.769 ± 40.716 | 40 | 1.318 ± 1.842 |
| 5 | 3.473 ± 2.752 | 20 | 1.569 ± 1.986 | 20 | 157.770 ± 256.236 | 20 | 1.678 ± 0.413 |
| 2.5 | 3.830 ± 0.652 | 10 | 1.560 ± 1.111 | 10 | 219.975 ± 373.969 | 10 | 16.543 ± 6.472 |
| 1.25 | 4.606 ± 4.259 | 5 | 2.257 ± 2.691 | 5 | 93.267 ± 151.878 | 5 | 13.524 ± 17.627 |
| 0.63 | 4.137 ± 3.685 | 2.5 | 2.852 ± 3.049 | 2.5 | 70.043 ± 1.2.809 | 2.5 | 12.181 ± 12.433 |
| 0.31 | 4.336 ± 4.548 | 1.25 | 1.370 ± 1.318 | 1.25 | 86.373 ± 132.169 | 1.25 | 5.056 ± 4.522 |

**CYP2B6**

| **Prochloraz** | | **Atrazine** | | **Pyrimethanil** | | **Chlorpyrifos-methyl** | |
| --- | --- | --- | --- | --- | --- | --- | --- |
| µg/ml | n-fold induction (±SD) | µg/ml | n-fold induction (±SD) | µg/ml | n-fold induction (±SD) | µg/ml | n-fold induction (±SD) |
| 10 | 0.376 ± 0.431 | 40 | 1.154 ± 1.038 | 40 | 2.437 ± 3.477 | 40 | 1.937 ± 2.446 |
| 5 | 0.549 ± 0.524 | 20 | 1.809 ± 1.999 | 20 | 5.141 ± 8.419 | 20 | 2.624 ± 3.169 |
| 2.5 | 0.399 ± 0.139 | 10 | 2.683 ± 0.518 | 10 | 129.923 ± 224.029 | 10 | 59.568 ± 91.734 |
| 1.25 | 0.791 ± 0.635 | 5 | 2.842 ± 2.209 | 5 | 288.522 ± 499.132 | 5 | 47.874 ± 80.955 |
| 0.63 | 1.234 ± 0.890 | 2.5 | 3.530 ± 2.168 | 2.5 | 412.502 ± 713.707 | 2.5 | 41.903 ± 72.356 |
| 0.31 | 1.636 ± 1.445 | 1.25 | 1.426 ± 1.041 | 1.25 | 44.554 ± 76.345 | 1.25 | 15.146 ± 25.734 |

**CYP3A4**

| **Prochloraz** | | **Atrazine** | | **Pyrimethanil** | | **Chlorpyrifos-methyl** | |
| --- | --- | --- | --- | --- | --- | --- | --- |
| µg/ml | n-fold induction (±SD) | µg/ml | n-fold induction (±SD) | µg/ml | n-fold induction (±SD) | µg/ml | n-fold induction (±SD) |
| 10 | 1.041 ± 0.937 | 40 | 1.688 ± 1.352 | 40 | 1.971 ± 1.770 | 40 | 2.934 ± 2.825 |
| 5 | 1.589 ± 0.680 | 20 | 1.324 ± 1.212 | 20 | 1.647 ± 0.891 | 20 | 8.169 ± 11.910 |
| 2.5 | 1.583 ± 0.802 | 10 | 1.553 ± 0.054 | 10 | 2.621 ± 1.830 | 10 | 6.620 ± 8.761 |
| 1.25 | 0.877 ± 0.306 | 5 | 1.422 ± 0.678 | 5 | 0.839 ± 0.484 | 5 | 13.939 ± 21.593 |
| 0.63 | 1.297 ± 1.169 | 2.5 | 2.097 ± 1.083 | 2.5 | 0.264 ± 0.064 | 2.5 | 2.129 ± 3.319 |
| 0.31 | 1.068 ± 0.913 | 1.25 | 0.526 ± 0.446 | 1.25 | 0.506 ± 0.663 | 1.25 | 3.332 ± 4.989 |

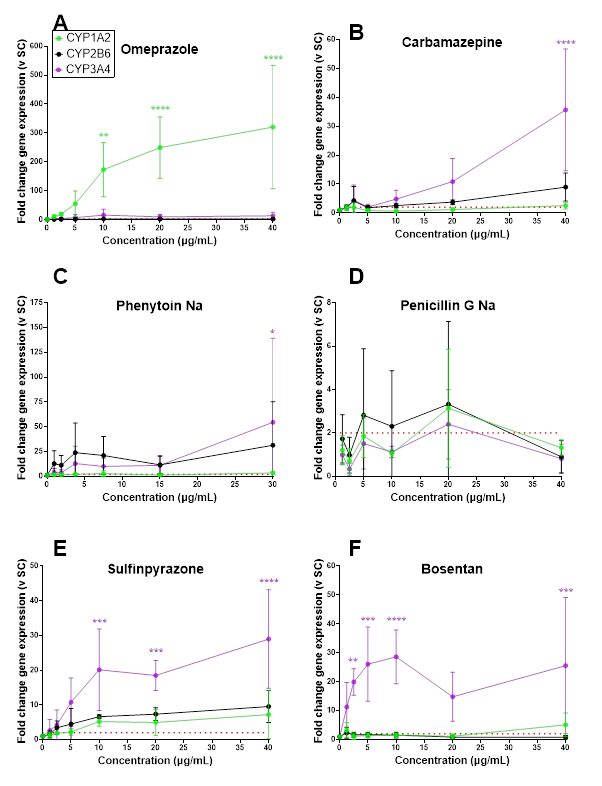

**Fig. S3A-F: Concentration response gene expression for augmentation chemicals tested.**

CYP1A2 (green), CYP2B6 (black) and CYP3A4 (purple) are shown. (A) Omeprazole; (B) Carbamazepine; (C) Phenytoin Sodium; (D) Penicillin G Sodium; (E) Sulfinpyrazone; (F) Bosentan hydrate. Three independent replicates (n = 3) (mean ± SD). * = P ≤ 0.05; ** = P ≤ 0.01; *** = P ≤ 0.001; **** = P ≤ 0.0001. GraphPad Prism Software v11.

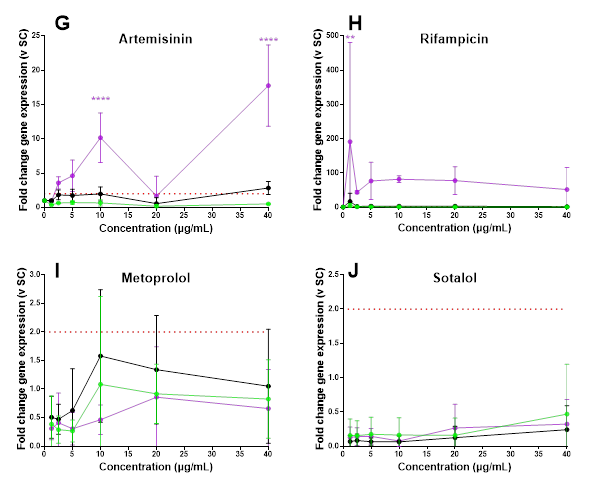

**Fig. S3G-J: Concentration response gene expression for augmentation chemicals tested.**

CYP1A2 (green), CYP2B6 (black) and CYP3A4 (purple) are shown. (G) Artemisinin; (H) Rifampicin; (I) Metoprolol; (J)Sotalol hydrochloride. Three independent replicates (n = 3) (mean ± SD). ** = P ≤ 0.01;; **** = P ≤ 0.0001. GraphPad Prism Software v11.

**Tab. S8. Statistical CYP mRNA induction data for all chemicals**.

Statistical significance of cell batches for CYP1A2, CYP2B6 and CYP3A4 mRNA induction. A one sample t-test compared against the solvent control. Green-shading indicates significance. Red indicates significant decrease. ns = P > 0.05; * = P ≤ 0.05; ** = P ≤ 0.01; *** = P ≤ 0.001; **** = P ≤ 0.0001; **** = p<0.0001P = inducer; N = non-inducer; N/A = insufficent prediction. Ome = Omeprazole; Carb = Carbamazepine; Phen Na = Phenytoin Sodium; Bos = Bosentan; Sot = Sotalol; Rif = Rifampicin; Met = Metoprolol; Art = Artemisinin; Sulf = Sulfinpyrazone; Teb = Tebuconazole; Benf = Benfuracarb; Chlor = Chlorpyrifos; DEET = N,N-diethyl-m-toluamide; Fip = Fipronil; Perm = Permethrin; Pro = Prochloraz; Atra = Atrazine; Pyrim = Pyrimethanil; Chlor-m = Chlorpyrifos-methyl

| **CYP1A2** | | | | | | | | | | | | | | | | | | | | |
| --- | --- | --- | --- | --- | --- | --- | --- | --- | --- | --- | --- | --- | --- | --- | --- | --- | --- | --- | --- | --- |
| **Result** | **P** | **B** | **B** | **N** | **P** | **P** | **N** | **P** | **N** | **N** | **P** | **P** | **P** | **N** | **N** | **E** | **P** | **E** | **P** | **N** |
|  | **Ome** | **Carb** | **Phen Na** | **Pen G Na** | **Sulf** | **Bos** | **Art** | **Rif** | **Met** | **Sot** | **Teb** | **Benf** | **Chlor** | **DEET** | **Fip** | **Perm** | **Pro** | **Atra** | **Pyrim** | **Chlor-m** |
| **Exp 1** | ns | ns | ns | ns | ns | ns | ** ↓ | ns | ****↓ | ****↓ | ns | * | ns | ** ↓ | ns | ns | ns | * | ns | ns |
| **Exp 2** | * | ns | ns | ns | ns | ns | ns | ns | ns | * ↓ | ns | ns | ns | ns | ns | ns | ns | ns | ns | ns |
| **Exp 3** | * | ns | ns | ns | ns | * ↓ | ** ↓ | ns | ns | ****↓ | ns | * | ** | ns | ns | ns | ns | ns | * | ns |
| **CYP2B6** | | | | | | | | | | | | | | | | | | | | |
| **Result** | **P** | **P** | **P** | **E** | **P** | **E** | **E** | **P** | **E** | **N** | **P** | **P** | **N** | **P** | **E** | **E** | **P** | **P** | **P** | **E** |
|  | **Ome** | **Carb** | **Phen Na** | **Pen G Na** | **Sulf** | **Bos** | **Art** | **Rif** | **Met** | **Sot** | **Teb** | **Benf** | **Chlor** | **DEET** | **Fip** | **Perm** | **Pro** | **Atra** | **Pyrim** | **Chlor-m** |
| **Exp 1** | ns | * | * | ns | ns | * | ns | ns | ns | ****↓ | * | * | ns | ns | ns | ns | ns | ** | ** | ns |
| **Exp 2** | ns | ** | * | ns | * | ns | ns | * | ns | ***↓ | ns | ** | ns | * | ns | ns | ns | ns | ns | ns |
| **Exp 3** | ns | ns | ** | ns | * | ns | * | ns | ns | ****↓ | ns | * | ns | ns | * | ** ↓ | ns | ns | * | ns |
| **CYP3A4** | | | | | | | | | | | | | | | | | | | | |
| **Result** | **P** | **P** | **P** | **E** | **P** | **P** | **P** | **P** | **N** | **N** | **P** | **P** | **P** | **P** | **P** | **N** | **P** | **P** | **P** | **E** |
|  | **Ome** | **Carb** | **Phen Na** | **Pen G Na** | **Sulf** | **Bos** | **Art** | **Rif** | **Met** | **Sot** | **Teb** | **Benf** | **Chlor** | **DEET** | **Fip** | **Perm** | **Pro** | **Atra** | **Pyrim** | **Chlor-m** |
| **Exp 1** | ns | * | ns | ns | * | * | ns | ns | **** ↓ | ****↓ | * | * | ns | * | * | ns | ns | ns | ns | ns |
| **Exp 2** | ns | * | * | ns | ns | ** | ns | * | ns | ** ↓ | ns | ns | ns | * | * | ns | ns | ns | * | ns |
| **Exp 3** | * | ns | * | ns | ns | ** | ns | * | ** ↓ | ****↓ | * | ns | * ↓ | * | * | ns | ns | ns | * | ns |
